# Brain-wide mapping of contextual fear memory engram ensembles supports the dispersed engram complex hypothesis

**DOI:** 10.1101/668483

**Authors:** Dheeraj S. Roy, Young-Gyun Park, Sachie K. Ogawa, Jae H. Cho, Heejin Choi, Lee Kamensky, Jared Martin, Kwanghun Chung, Susumu Tonegawa

## Abstract

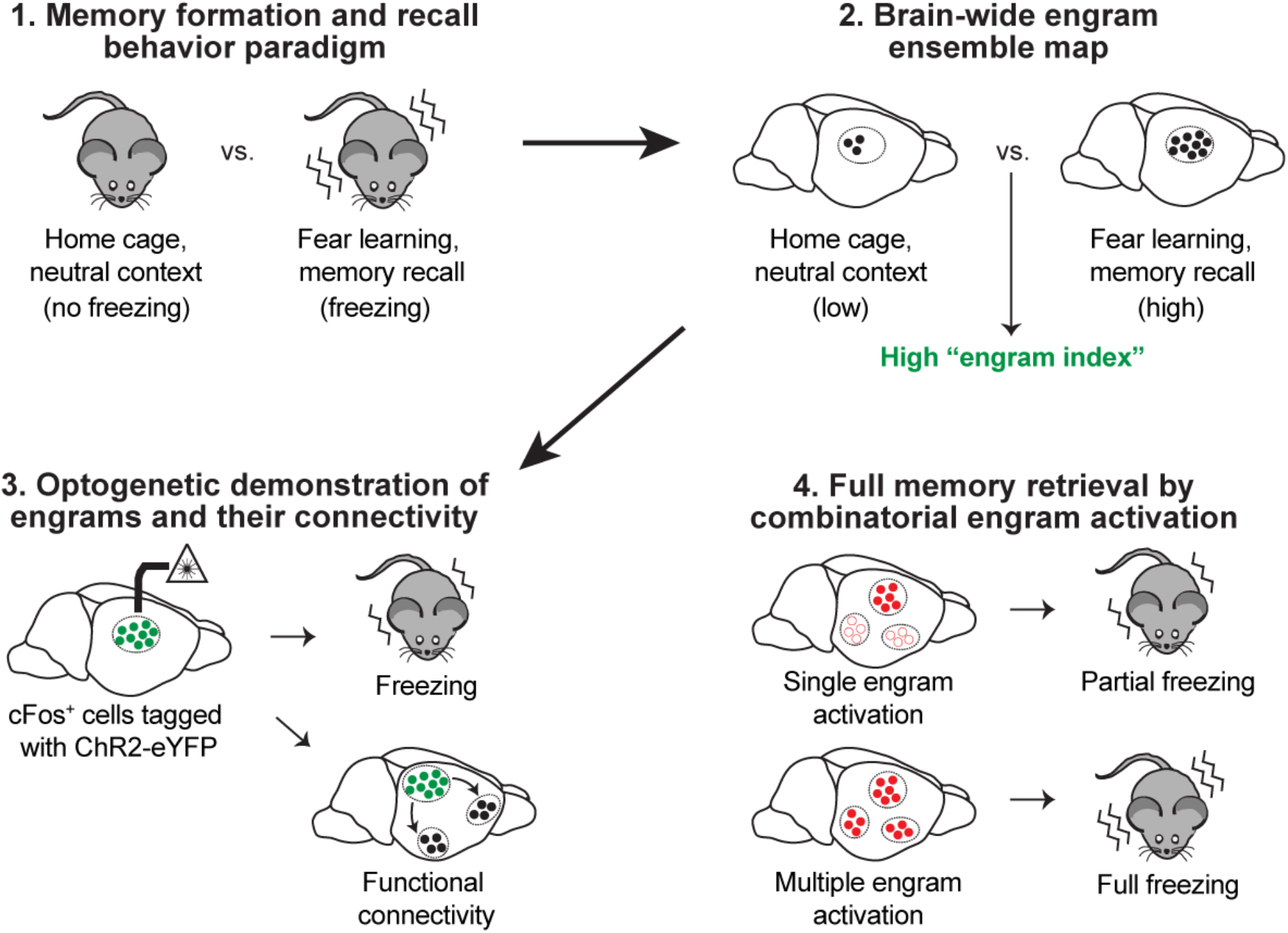

**SUMMARY:** Neuronal ensembles that hold specific memory (memory engrams) have been identified in the hippocampus, amygdala, and cortex. It has been hypothesized that engrams for a specific memory are distributed among multiple brain regions that are functionally connected. Here, we report the hitherto most extensive engram map for contextual fear memory by characterizing activity-tagged neurons in 409 regions using SHIELD-based tissue phenotyping. The mapping was aided by a novel engram index, which identified cFos^+^ brain regions holding engrams with a high probability. Optogenetic manipulations confirmed previously known engrams and revealed new engrams. Many of these engram holding-regions were functionally connected to the CA1 or amygdala engrams. Simultaneous chemogenetic reactivation of multiple engrams, which mimics natural memory recall, conferred a greater level of memory recall than reactivation of a single engram ensemble. Overall, our study supports the hypothesis that a memory is stored in functionally connected engrams distributed across multiple brain regions.

## INTRODUCTION

A memory engram is the enduring physical or chemical changes that occur in brain networks upon learning, representing acquired memory information. Engrams are held by a set of neuronal ensembles that are activated by learning, and a reactivation of these neurons gives rise to recall of the specific memory (Josselyn et al., 2015; Semon, 1904; Tonegawa et al., 2015).

Several studies have identified engram cells for different memories in many brain regions including the hippocampus (Liu et al., 2012; Ohkawa et al., 2015; Roy et al., 2016), amygdala (Han et al., 2009; Redondo et al., 2014), retrosplenial cortex (Cowansage et al., 2014), and prefrontal cortex (Kitamura et al., 2017). In particular, by performing engram cell connectivity and engram cell reactivation experiments, it has been demonstrated that a given memory is stored in a specific set of engram cell ensembles that exhibit enduring cellular changes and are functionally connected (Roy et al., 2017a; Ryan et al., 2015). While these findings have enhanced our understanding of engrams and their dynamics, a thorough mapping of engram cell ensemble circuits for a specific memory has not been accomplished.

The development of tissue clearing techniques, such as CLARITY (Chung et al., 2013) and iDISCO (Renier et al., 2014), combined with advanced microscopy has enabled high-throughput analyses of intact mouse brain samples. Brain-wide activity mapping has been applied to parental behavior (Renier et al., 2016), foot shocks and cocaine administration (Ye et al., 2016), and fear conditioning and remote memory retrieval (DeNardo et al., 2019; Vetere et al., 2017). These studies, however, either did not ascertain whether the activated neurons were engram-holding ensembles or analyzed only a limited number of brain regions. To further our understanding of the organization of memory engram cell ensembles, brain-wide neuronal activity patterns following memory formation should be examined in a more holistic manner.

Importantly, since engram cell ensembles formed during learning are preferentially reactivated during recall of the specific memory (Josselyn et al., 2015; Semon, 1904; Tonegawa et al., 2015), comparing brain-wide neuronal activity maps following memory encoding and memory recall will help identify putative memory engram ensembles.

In the present study, we used these criteria to identify engram cell ensembles and their high probability candidates in a brain-wide manner, by following a three-step approach. First, we applied SHIELD-based brain-wide tissue phenotyping (Park et al., 2018) for mapping activity-tagged neurons in the Fos-TRAP mouse line (Guenthner et al., 2013). Using an unbiased and automated approach, we cross-compared the activity of 409 brain regions during contextual fear memory encoding and recent memory recall. Second, because to conduct optogenetic or chemogenetic manipulations that permit an assertion of engrams in hundreds of brain regions that displayed activity-dependent labeling is impractical, we devised an engram index to rank-order putative engram cell ensemble candidates (see STAR Methods). Third, focusing on a dozen brain regions that showed a high engram index value, we conducted optogenetic manipulations. This not only confirmed previously identified engrams but also revealed engrams in new brain regions. Many of these engrams and engram candidates were functionally connected to hippocampal CA1 and/or basolateral amygdala engrams. Finally, simultaneous reactivation of multiple engram ensembles using chemogenetics conferred greater levels of memory recall than reactivation of their subsets as would be expected from the natural process of memory recall. Together, our study supports the concept that a specific memory is stored in functionally connected engram cell ensembles that are widely distributed in multiple brain regions.

## RESULTS

### Integration of activity-dependent labeling with SHIELD tissue clearing

The first step of our unbiased engram mapping approach required brain-wide activity-dependent cell labeling. Expression of immediate-early genes (IEGs) has been used to visualize neuronal activity in a given brain region (Kubik et al., 2007). By linking the expression of Cre^ERT2^ to the IEG *c-fos* in a tamoxifen-dependent manner, the Fos-TRAP mouse line permits brain-wide labeling of activated neurons within a user-defined time window of several hours (Guenthner et al., 2013). We crossed Fos-TRAP mice with a Cre-dependent tdTomato reporter mouse line (Figure 1A). Using the fast-acting 4-hydroxytamoxifen (4-OHT), we prepared four behavioral cohorts: mice that received 4-OHT and remained in their home cage for the labeling duration (home cage group), mice that received 4-OHT followed by a context exposure and were returned to their home cage (context group), mice that received 4-OHT followed by contextual fear conditioning training and were returned to their home cage (CFC group), and mice that received 4-OHT followed by fear memory recall and were returned to their home cage (recall group). One week after labeling, brains were SHIELD-processed (Park et al., 2018), which preserves endogenous tdTomato fluorescence and tissue architecture during the clearing process. The optically transparent SHIELD brains were imaged using a custom-built high-speed selective plane illumination microscope (SPIM) (Figure S1, and see Movies S1 and S2). Due to the high quality of tissue architecture preservation during the clearing and imaging processes, three-dimensional (3D) brain images could be automatically aligned to a standard mouse brain atlas (see STAR Methods), and further, we could successfully perform automatic brain region segmentation. Crucially, by applying a neural network-based automatic cell counting algorithm, we detected tdTomato-positive activated cells in the entire brain sample at single cell resolution; representative examples from home cage and CFC groups are shown in Figure 1A. These experiments validated our brain-wide activity-dependent mapping strategy.

**Figure 1.**
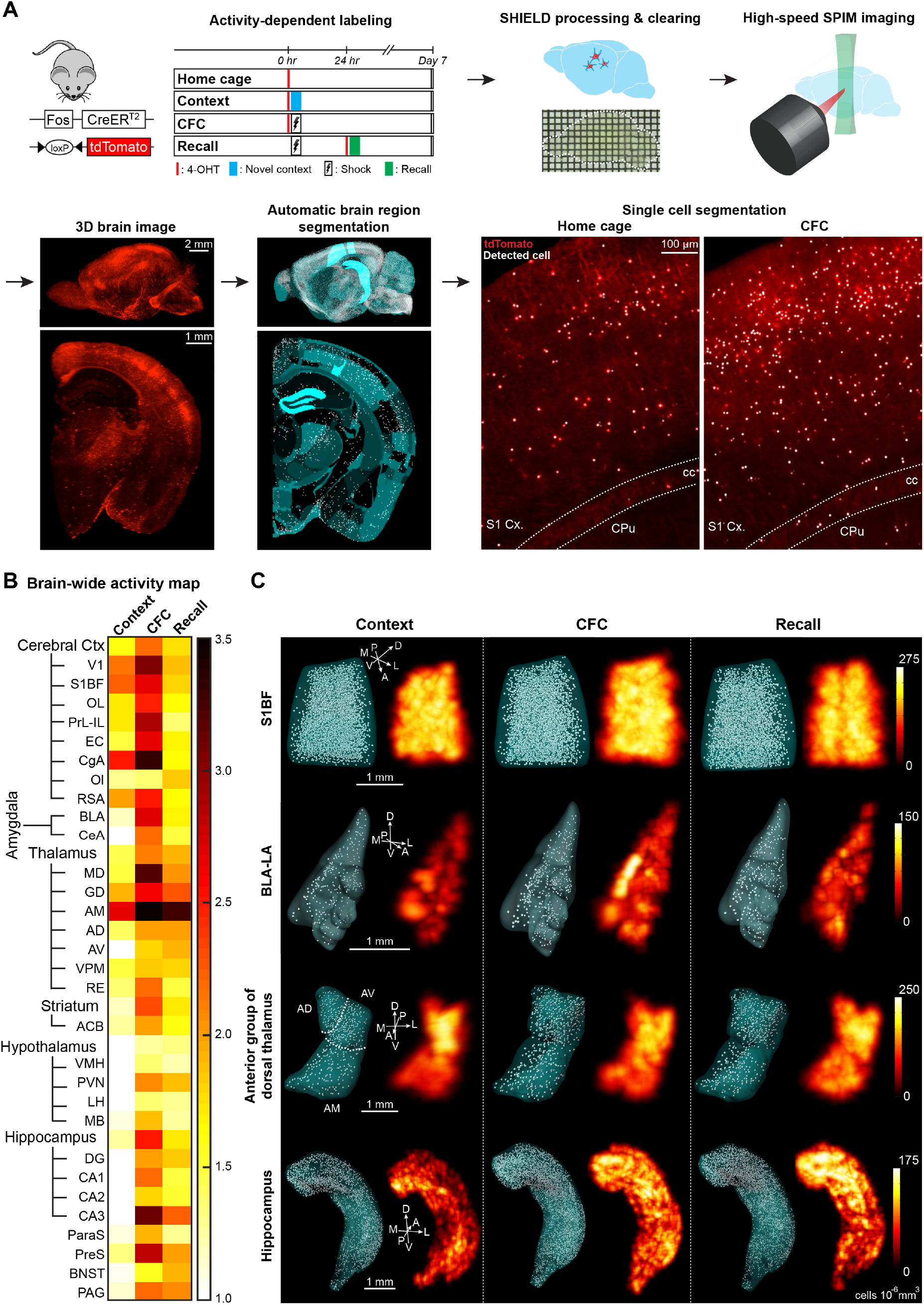
Generation of behavior epoch-specific brain-wide activity maps. (A) Activity mapping pipeline. We used Fos-TRAP mice crossed with a Cre-dependent tdTomato reporter mouse line to prepare four behavioral cohorts: home cage, context exposure (Context), contextual fear conditioning (CFC), and fear memory recall (Recall). Brain samples were used for SHIELD processing, SPIM imaging, 3D reconstructions, automatic brain region segmentation, and automatic single activated-cell detection. Primary somatosensory cortex (S1 Cx), corpus callosum (cc), caudate putamen (CPu). (B) Brief, representative version of our brain-wide activity mapping results (n = 4 mice each for home cage and CFC groups, n = 3 mice each for context and recall groups), where the behavioral groups are normalized to home cage data. Color-coded scale bar represents fold increases in the numbers of activated neurons relative to home cage. Primary visual cortex (V1), primary somatosensory cortex barrel field (S1BF), orbital cortex (OL), prelimbic cortex (PrL), infralimbic cortex (IL), entorhinal cortex (EC), anterior cingulate cortex (CgA), olfactory cortex (Ol), retrosplenial cortex (RSA), basolateral amygdala (BLA), central amygdala (CeA), mediodorsal thalamus (MD), dorsal geniculate thalamus (GD), anteromedial thalamus (AM), anterodorsal thalamus (AD), anteroventral thalamus (AV), ventroposterior medial nucleus of thalamus (VPM), nucleus reuniens of thalamus (RE), nucleus accumbens (ACB), ventromedial hypothalamus (VMH), paraventricular hypothalamus (PVN), lateral hypothalamus (LH), mammillary body (MB), hippocampal dentate gyrus (DG), hippocampal CA1 (CA1), hippocampal CA2 (CA2), hippocampal CA3 (CA3), para-subiculum (ParaS), pre-subiculum (PreS), bed nucleus of the stria terminalis (BNST), periaqueductal gray (PAG). For a full list of brain regions analyzed, refer to Table S1. (C) 3D rendering of four brain regions, specifically S1BF cortex, BLA-LA, anterior group of dorsal thalamus, and hippocampus, showing automatically segmented activated neuronal populations along with corresponding heat maps. Lateral amygdala (LA). Dorsal (D), ventral (V), anterior (A), posterior (P), medial (M), lateral (L).

### Generation of behavior epoch-specific brain-wide activity (cFos) maps

We quantified the number of activated neurons in 409 brain regions (see Table S1) from the home cage group, context exposure group, memory encoding group, and memory recall group. To generate brain-wide activity maps, the tdTomato (i.e., cFos^+^) cell counts of context, CFC encoding, and CFC recall groups were normalized by that of the home cage group (Figure 1B). In the representative brain-wide activity map (Figure 1B), we plotted the activation pattern for entire brain structures including cerebral cortex, amygdala, thalamus, striatum, hypothalamus, hippocampus, and their respective sub-divisions. At the brain-wide level, it is clear that CFC encoding results in the activation of many brain regions as compared to exposure to a neutral valence context. On the other hand, while memory recall activated fewer neurons as compared to memory encoding, brain regions that were most strongly activated by recall were those that were also activated by encoding. This observation is consistent with the notion that memory engram ensembles that are activated by learning are reactivated during recall of the specific memory (Semon, 1904; Tonegawa et al., 2015).

Among the most robustly activated brain regions by memory encoding were the prefrontal cortex, entorhinal cortex (EC), amygdala, hippocampus, and periaqueductal gray (PAG) (Figures 1B and 1C). Importantly, as it has been reported that these structures play a crucial role in contextual fear learning and memory (Tovote et al., 2015), this activation pattern supports the accuracy of our brain-wide activity-mapping results. We also observed that sensory cortical regions, specifically primary visual cortex (V1) and primary somatosensory cortex barrel field (S1BF), showed minimal differences between context exposure and CFC memory encoding epochs, suggesting that these structures are less likely to contain engram cell populations for this memory. Other than learning-activated brain regions and sensory brain regions, a third type of activity pattern was observed in hypothalamic sub-regions, which showed weaker but significant overall activation in this behavioral paradigm as compared to the above-mentioned learning-activated and sensory brain structures. Nevertheless, two hypothalamic nuclei, paraventricular hypothalamus (PVN) and mammillary bodies (MB) were activated by CFC encoding, and further the PVN was also activated by memory recall.

In these brain-wide activity maps, we found several regions where the neurons were strongly activated by both encoding and recall, but their role in memory formation has previously been less understood as compared to hippocampal and amygdala regions (Figure 1C). These regions include thalamic structures, namely the anteromedial thalamus (AM) sub-division of the anterior group of dorsal thalamus, mediodorsal thalamus (MD), and nucleus reuniens thalamus (RE), as well as the striatum, para-subiculum (ParaS), and pre-subiculum (PreS). In order to narrow down the list of putative engram-containing brain regions, we created an engram index (see STAR Methods) and ordered brain regions that displayed higher cFos activation levels in both CFC encoding and recall epochs, as compared to both home cage and context exposure epochs (Figure 2, see Figure S2 and Table S2). This index provides an unbiased criterion to identify brain regions that potentially carry memory engram cells (see Movie S3). Hippocampal, amygdala, and association cortical brain regions showed high engram index values (Figure 2), which supports the validity of this approach.

**Figure 2.**
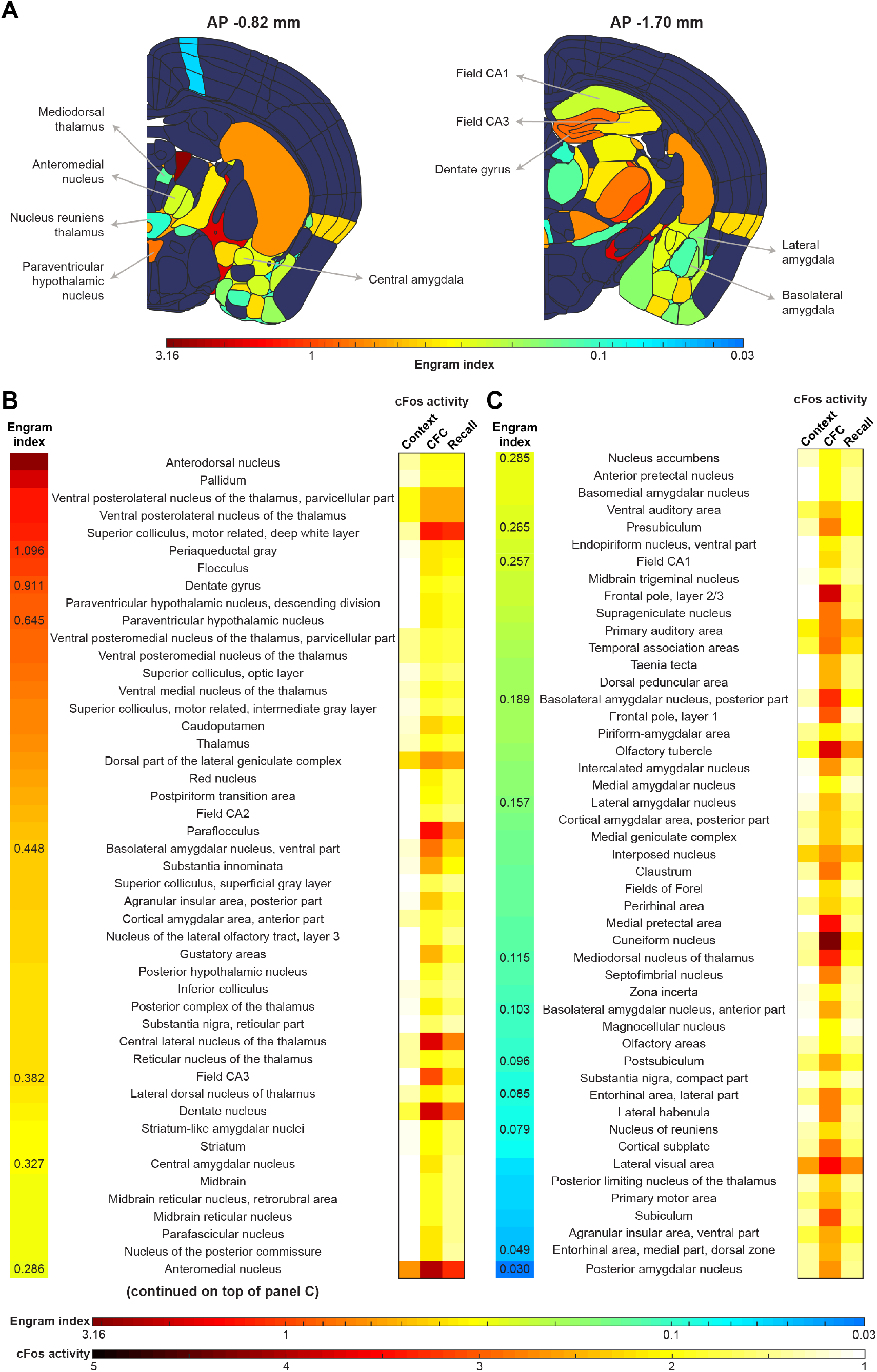
Brain-wide engram indices. (A) Representative coronal brain section outlines showing color-coded memory engram indices (see STAR Methods) across brain structures. Brain regions with significant engram indices are colored, whereas other brain regions are shown in dark blue shading. Distance from bregma along the anterior-posterior (AP) axis has been indicated for each coronal section. (B and C) Rank-ordered list of 97 brain regions with engram indices greater than 0. Engram index and cFos activation during behavioral epochs are color-coded and plotted (n = 4 mice each for home cage and CFC groups, n = 3 mice each for context and recall groups). High engram index brain regions are those with greatest neuronal activity during both memory encoding and recall epochs (relative to both home cage and context groups). For a full list of significant brain regions, including individual layers within regions, and their corresponding engram indices, refer to Table S2.

An advantage of using intact tissue clearing rather than traditional brain sectioning-based histology for activity-dependent labeling studies is that in cleared brain samples we can quantitatively characterize 3D activity patterns across the entirety of the brain without sampling errors (see Movies S1-S3). Representative examples of 3D rendering accompanied by activity heat maps of our brain-wide data (Figure 1C) makes it possible to determine whether neuronal activity changes in a specific brain region are restricted to the dorsal-ventral, anterior-posterior, or medial-lateral axes. By quantifying intra-regional activation patterns, we observed that dorsal hippocampus showed significantly greater activation during CFC encoding as compared to both home cage and context exposure groups, which was not the case in ventral hippocampus although we did observe a trend in a similar direction (Figures 3A-3C). This pattern is consistent with previous studies on the crucial role of dorsal hippocampus in contextual learning (Fanselow and Dong, 2010). Recently, it was reported that anterior basolateral amygdala (BLA) contain *Rspo2*-positive excitatory neurons that underlie negative-valence stimuli-induced behaviors such as fear learning (Kim et al., 2016). Consistent with this report, we observed greater activation of anterior BLA during CFC encoding as compared to posterior BLA (Figures 3A, 3B, and 3D). Given the known important role of EC circuits in memory formation, we observed differential inter-group activation patterns in medial EC (MEC) vs. lateral EC (LEC) along their respective medial-lateral axes, suggesting that the size of engram ensembles may vary along this structure (Figures 3A, 3B, 3E, and 3F). Finally, we examined the 3D distribution pattern of activated neurons in the thalamic reticular nucleus (TRN) (Figures 3A, 3B, and 3G). Interestingly, although the TRN consists mostly of GABAergic neurons, we found greater neuronal activation during context exposure and CFC encoding in dorsal TRN as compared to ventral TRN, which may reflect an enhanced contribution of dorsal TRN circuitry to the suggested cognitive functions of this brain structure (Pinault, 2004).

**Figure 3.**
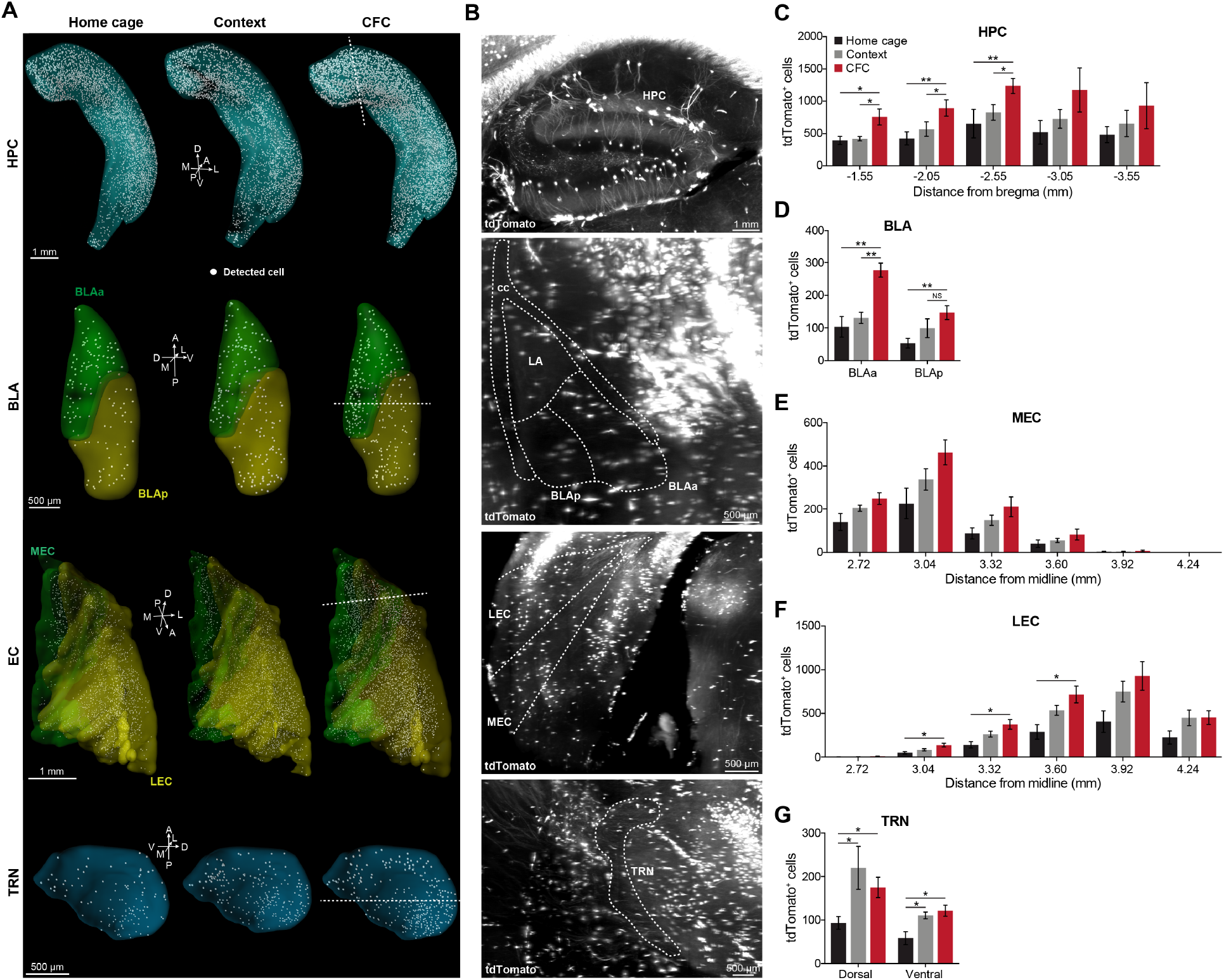
Intra-regional patterns of activated neuronal ensembles across axes. (A) Representative 3D rendering of hippocampus (HPC), BLA, EC, and thalamic reticular nucleus (TRN). BLA rendering distinguishes anterior vs. posterior sub-regions. EC rendering distinguishes medial vs. lateral sub-regions. Green shading represents anterior BLA (BLAa) or medial EC (MEC). Yellow shading represents posterior BLA (BLAp) or lateral EC (LEC). (B) tdTomato images showing activated neurons corresponding to the optical sections indicated by the white dotted lines in Figure 3A. (C-G) Quantification of the intra-regional distribution of activated (tdTomato^+^) neuronal ensembles (n = 4 mice each for home cage and CFC groups, n = 3 mice for context), in HPC (C), BLA (D), MEC (E), LEC (F), and TRN (G). Anterior-posterior axis for HPC and BLA. Medial-lateral axis for MEC and LEC. Dorsal-ventral axis for TRN. Millimeter (mm). NS, not significant. Statistical comparisons are performed using a one-way ANOVA followed by Tukey multiple comparison post-hoc tests; *P < 0.05, **P < 0.01. Data are presented as mean ± SEM.

### Stimulation frequency-dependent memory retrieval by optogenetic engram cell reactivation

Although the initial demonstration showed that optogenetic reactivation of hippocampal dentate gyrus engram cells at a frequency of 20 Hz resulted in recall of the specific memory (Liu et al., 2012), following studies showed that engram cell ensembles in hippocampal CA1 (Ryan et al., 2015) and prefrontal cortex (Kitamura et al., 2017) are more effectively reactivated using a 4 Hz optogenetic frequency protocol rather than a 20 Hz protocol. These findings indicate that the identification of novel engram cell ensembles must try multiple frequencies of optogenetic stimulation since certain brain regions may prefer a specific frequency range. Therefore, in this study we performed optogenetic reactivation at both 4 Hz and 20 Hz. To tag putative engram cells, we employed a double virus approach (Roy et al., 2016), which included an activity-dependent vector c-Fos-tTA and a channelrhodopsin-2 (ChR2) tagging vector TRE-ChR2-eYFP (Figure 4A). Neurons activated during CFC training were tagged by taking the animals off their doxycycline diet for 24 hours before the encoding epoch (Figure 4B). Following encoding, we performed a natural memory recall test to confirm successful fear memory recall, after which in a neutral context we reactivated putative engram cells at the two different frequencies (Figure 4B).

**Figure 4.**
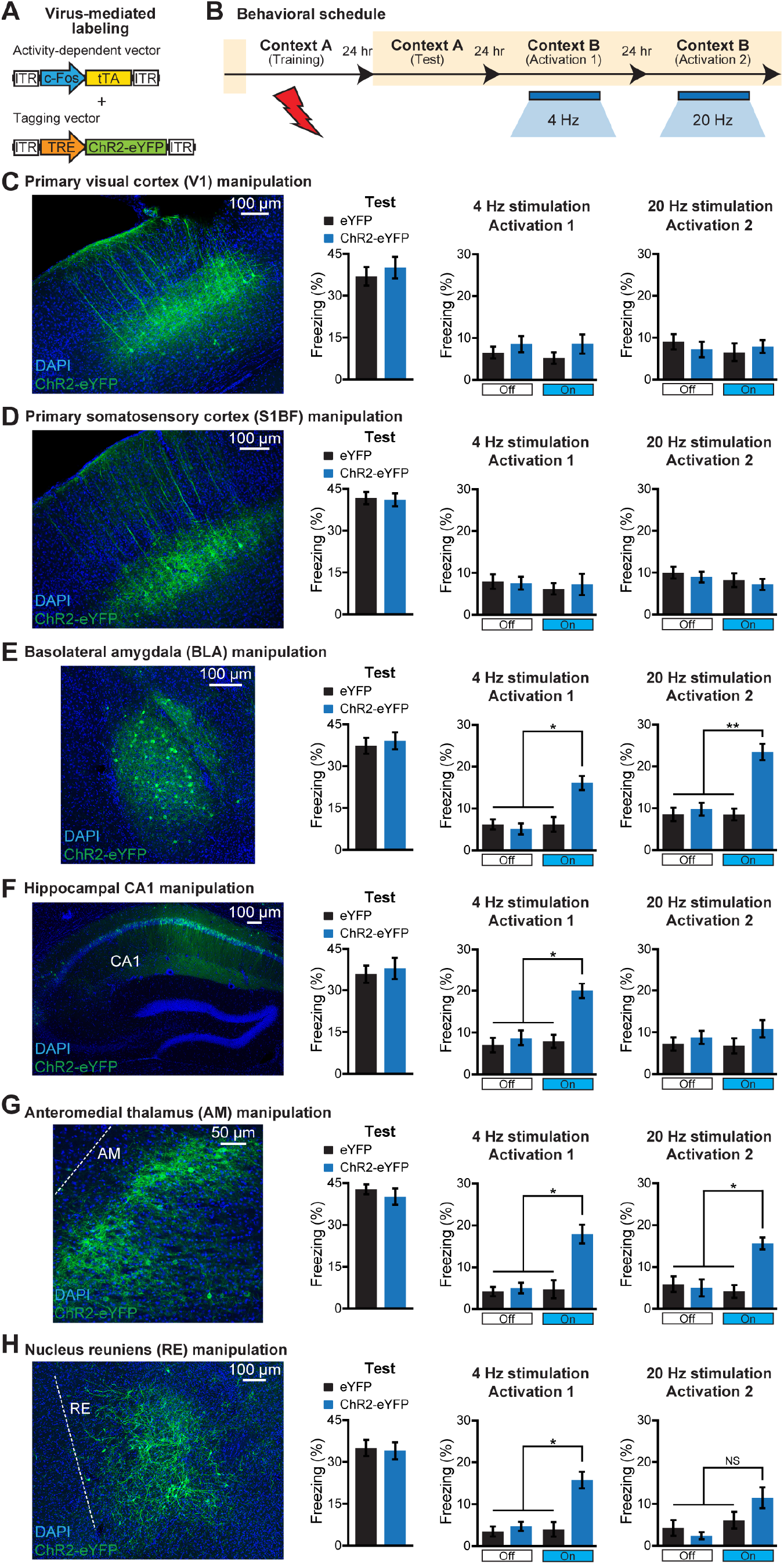
Stimulation frequency-dependent memory retrieval by activating engram cells. (A) Virus-mediated cFos^+^ neuronal labeling strategy using a cocktail of c-Fos-tTA and TRE-ChR2-eYFP. Wild-type mice raised on doxycycline (DOX) food were injected with the two viruses bilaterally in the respective target regions. (B) Behavioral schedule. Beige shading signifies that mice were on a DOX diet, precluding ChR2-eYFP expression. Mice were taken off DOX 24 hrs before contextual fear conditioning (CFC). One day after training, a natural recall test was performed (Test). The next day, mice received an optogenetic reactivation (Activation 1) session in a neutral context B, using a 4 Hz stimulation protocol. The following day mice received a second optogenetic reactivation (Activation 2) session, this time using a 20 Hz stimulation protocol. (C) Primary visual cortex (V1) section showing cFos^+^ neurons labeled with ChR2-eYFP using the two-virus approach (left). Natural memory recall (Test). Optogenetic reactivation (Activation 1, Activation 2). eYFP (n = 7 mice) and ChR2-eYFP (n = 8 mice) groups. (D) Primary somatosensory cortex (S1BF) section showing cFos^+^ neurons labeled with ChR2-eYFP using the two-virus approach (left). Natural memory recall (Test). Optogenetic reactivation (Activation 1, Activation 2). eYFP (n = 8 mice) and ChR2-eYFP (n = 8 mice) groups. (E) Basolateral amygdala (BLA) section showing cFos^+^ neurons labeled with ChR2-eYFP using the two-virus approach (left). Natural memory recall (Test). Optogenetic reactivation (Activation 1, Activation 2). eYFP (n = 9 mice) and ChR2-eYFP (n = 10 mice) groups. (F) Hippocampal CA1 section showing cFos^+^ neurons labeled with ChR2-eYFP using the two-virus approach (left). Natural memory recall (Test). Optogenetic reactivation (Activation 1, Activation 2). eYFP (n = 7 mice) and ChR2-eYFP (n = 7 mice) groups. (G) Anteromedial thalamus (AM) section showing cFos^+^ neurons labeled with ChR2-eYFP using the two-virus approach (left). Natural memory recall (Test). Optogenetic reactivation (Activation 1, Activation 2). eYFP (n = 10 mice) and ChR2-eYFP (n = 9 mice) groups. (H) Nucleus reuniens (RE) section showing cFos^+^ neurons labeled with ChR2-eYFP using the two-virus approach (left). Natural memory recall (Test). Optogenetic reactivation (Activation 1, Activation 2). eYFP (n = 11 mice) and ChR2-eYFP (n = 11 mice) groups. NS, not significant. Statistical comparisons are performed using unpaired *t* tests; *P < 0.05, **P < 0.01. Data are presented as mean ± SEM.

Based on our brain-wide neuronal activity map (Figure 1B, and see Figure S2), it is clear that cortical regions V1 and S1BF exhibit increased activity during CFC encoding with some activation during recall. However, we were unable to induce memory recall by optogenetic reactivation of either cortical region (Figures 4C and 4D). We cannot rule out the possibility that these regions contain CFC engram cells, although activity differences between context exposure and CFC encoding and recall were relatively small (Figure 1B). As a positive control, we observed robust memory recall by the optogenetic reactivation of BLA engram cells at both 4 and 20 Hz frequencies (Figure 4E), as well as by the reactivation of dorsal CA1 engram cells at 4 Hz specifically (Figure 4F).

In contrast to sensory cortical regions, our brain-wide activity map and engram indices revealed several thalamic nuclei that were activated significantly by CFC memory encoding and recall, and less by context exposure. Since the contributions of thalamic nuclei to memory engrams remain understudied, we decided to focus on the thalamus and identified two structures from our engram indices that have been previously linked to spatial learning and navigation, the anterior group of dorsal thalamus (de Lima et al., 2017) and nucleus reuniens (RE) (Ramanathan et al., 2018). Within the anterior group of dorsal thalamus, we focused on anteromedial thalamus (AM), which showed robust neural activity during fear learning and memory recall, had a significant engram index value (Figure 2B), and could be effectively targeted using our viral approach. Strikingly, optogenetic reactivation of AM cells that were activated during fear learning resulted in memory recall at both 4 Hz and 20 Hz (Figure 4G). Similarly, reactivation of putative RE thalamus engram cells resulted in memory recall, but only at the 4 Hz frequency protocol (Figure 4H). Together, the application of brain-wide activity mapping followed by causal optogenetic reactivation experiments allowed us to identify two novel brain regions, AM thalamus and RE thalamus, that carry engram cells for CFC memory.

### Brain-wide neuronal activity patterns induced by optogenetic manipulations of engram cell ensembles

Despite recent advances in memory engram research, neuronal activity changes in downstream brain regions induced by the optogenetic reactivation of an upstream memory engram cell population remains unclear. To address this question, we examined cFos activation in a brain-wide manner including downstream brain regions following engram cell reactivation or inhibition of an upstream engram ensemble in the CFC paradigm. In the first set of experiments, we focused on hippocampal dorsal CA1 engram manipulations using the double virus approach (Figure 5A). Optogenetic reactivation of CA1 engram cells at 4 Hz resulted in robust memory recall, which was not observed in control eYFP mice (Figures 5A-5C). One hour after optogenetic reactivation of CA1 engram cells in the neutral context, mice were processed for cFos staining. Among 15 brain regions examined (Figure 5D, and see Figure S3A), the numbers of cFos-positive cells were most significantly increased in lateral hypothalamus (LH) and BLA in the CA1 ChR2-eYFP group as compared to the control group in which CA1 engram cells were not optogenetically activated (Figure 5I). EC and hippocampal dentate gyrus (DG) showed a pattern of increased activation. To perform a comparable analysis following CA1 engram cell inhibition, we employed a double virus approach expressing eArchT-mCherry in engram cells by injection of an activity-dependent vector c-Fos-tTA and an enhanced archaerhodopsin (eArchT) tagging vector TRE-eArchT-mCherry. Optogenetic inhibition of CA1 engram cells decreased memory recall in the training context as compared to control mCherry-only mice (Figures 5E-5G). cFos analyses following CA1 engram cell inhibition (Figure 5H, and see Figure S3B) revealed that prefrontal sub-regions, infralimbic (IL) and prelimbic (PrL) cortices, had significantly decreased neuronal activity as compared to the control group (Figure 5I). Further, nucleus accumbens shell (AcbSh), BLA, central amygdala (CeA), DG, and PAG showed a pattern of decreased activation.

**Figure 5.**
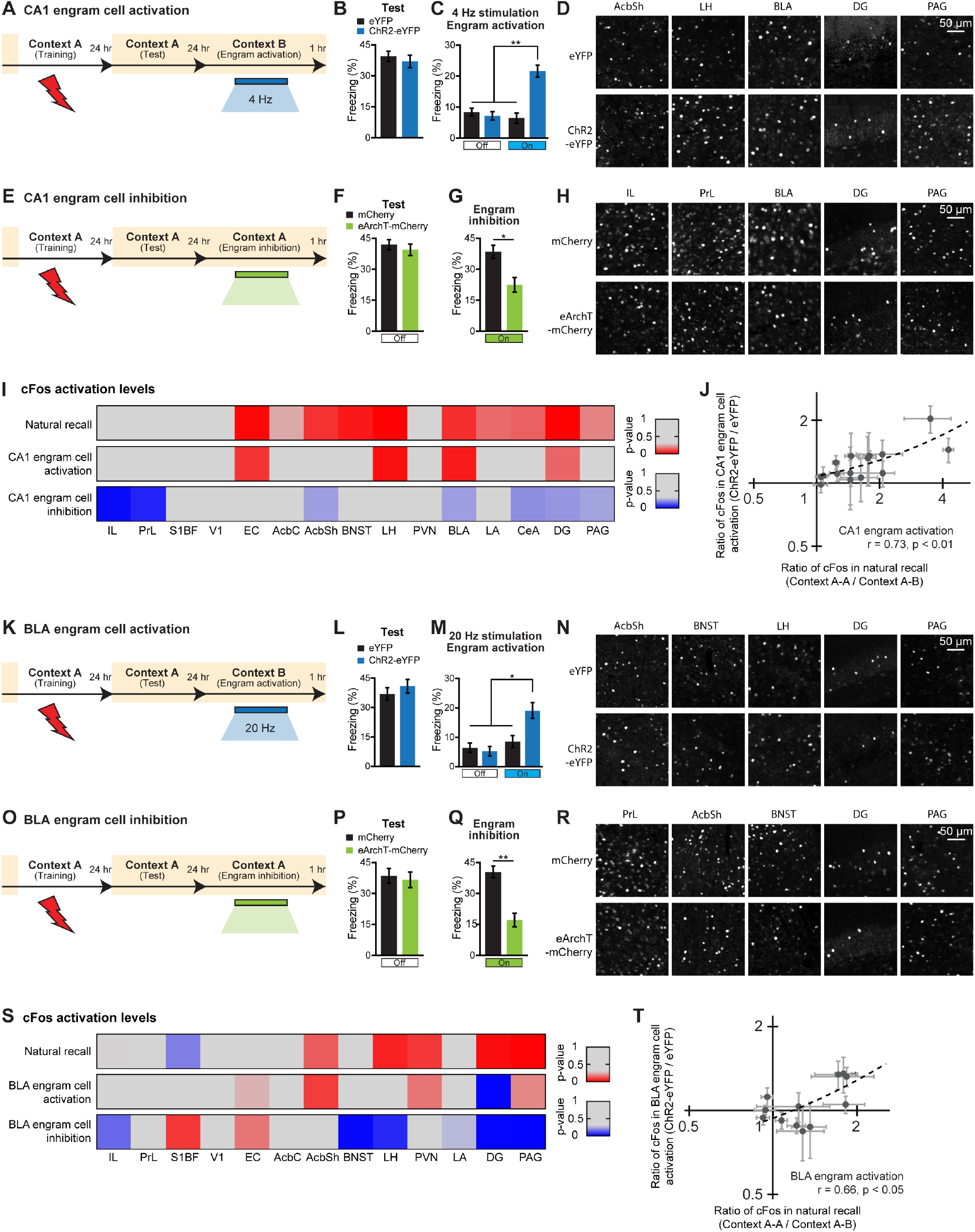
Brain-wide neuronal activity patterns following engram cell manipulations. (A) Behavioral schedule. Beige shading signifies that mice were on a DOX diet, precluding ChR2-eYFP expression. Mice were taken off DOX 24 hrs before contextual fear conditioning (CFC). One day after training, a natural recall test was performed (Test). The next day, mice received an optogenetic reactivation (Engram activation) session in a neutral context B, using a 4 Hz stimulation protocol. (B) Natural memory recall (Test). (C) Optogenetic reactivation (Engram activation). eYFP (n = 8 mice) and ChR2-eYFP (n = 8 mice) groups. (D) cFos^+^ neurons in representative brain regions, including nucleus accumbens shell (AcbSh), LH, BLA, hippocampal DG, and PAG, shown for eYFP (top row) and ChR2-eYFP (bottom row) groups. (E) Behavioral schedule for CA1 engram cell inhibition. One day after training, a natural recall test was performed (Test). The next day, mice received an optogenetic inhibition (Engram inhibition) session in the training context. (F) Natural memory recall (Test). (G) Optogenetic inhibition (Engram inhibition). mCherry (n = 6 mice) and eArchT-mCherry (n = 8 mice) groups. (H) cFos^+^ neurons in representative brain regions, including IL, PrL, BLA, hippocampal DG, and PAG, shown for mCherry (top row) and eArchT-mCherry (bottom row) groups. (I) Heat map represents cFos activation levels in 15 brain regions for natural memory recall and CA1 engram cell manipulations (n = 3-4 mice per group). Red colored regions indicate an increase in the number of cFos^+^ neurons based on the *p*-value obtained by comparing control vs. manipulation group data, whereas blue colored regions indicate a decrease in the number of cFos^+^ neurons. (J) Scatter plot comparing the ratio of cFos^+^ cell counts between CA1 engram cell activation and natural memory recall (n = 3-4 mice per group). Pearson’s correlation coefficient (r). (K) Behavioral schedule for BLA engram cell activation, as performed in Figure 5A. Optogenetic reactivation (Engram activation) session used a 20 Hz stimulation protocol. (L) Natural memory recall (Test). (M) Optogenetic reactivation (Engram activation). eYFP (n = 7 mice) and ChR2-eYFP (n = 7 mice) groups. (N) cFos^+^ neurons in representative brain regions, including AcbSh, BNST, LH, hippocampal DG, and PAG, shown for eYFP (top row) and ChR2-eYFP (bottom row) groups. (O) Behavioral schedule for BLA engram cell inhibition. (P) Natural memory recall (Test). (Q) Optogenetic inhibition (Engram inhibition). mCherry (n = 7 mice) and eArchT-mCherry (n = 9 mice) groups. (R) cFos^+^ neurons in representative brain regions, including PrL, AcbSh, BNST, hippocampal DG, and PAG, shown for mCherry (top row) and eArchT-mCherry (bottom row) groups. (S) Heat map represents cFos activation levels in 13 brain regions for natural memory recall and BLA engram cell manipulations (n = 4 mice per group). Red colored regions indicate an increase in the number of cFos^+^ neurons based on the *p*-value obtained by comparing control vs. manipulation group data, whereas blue colored regions indicate a decrease in the number of cFos^+^ neurons. (T) Scatter plot comparing the ratio of cFos^+^ cell counts between BLA engram cell activation and natural memory recall (n = 4 mice per group). Pearson’s correlation coefficient (r). Unless specified, statistical comparisons are performed using unpaired *t* tests; *P < 0.05, **P < 0.01. Data are presented as mean ± SEM.

In the second set of experiments, we focused on optogenetic manipulations of BLA engram cells also using the double virus approach. As expected, BLA engram cell reactivation induced memory recall (Figures 5K-5M), and BLA engram cell inhibition decreased memory retrieval in the training context (Figures 5O-5Q). cFos analyses following optogenetic activation of BLA engram cells (Figure 5N, 5S, and see Figure S3C) showed a pattern of increased activation in EC, AcbSh, paraventricular hypothalamus (PVN), and PAG. Conversely, cFos analyses following BLA engram cell inhibition (Figure 5R, 5S, and see Figure S3D) showed significantly decreased activity in bed nucleus of the stria terminalis (BNST), DG, and PAG regions. Also, IL, LH, and lateral amygdala (LA) showed a pattern of decreased activation.

### Optogenetic manipulation mimics natural recall cue-driven effects in the engram ensemble pathway

The cFos analyses following optogenetic manipulations of engram cells allowed us to examine individual brain region changes in neuronal activity levels. Taking advantage of this data set, we wanted to examine the brain-wide pattern of neuronal activity induced by optogenetic manipulations of engram cells and compare them to natural memory recall activity patterns more directly. For this purpose, we plotted cFos activity levels for natural recall, ChR2-based engram reactivation groups, and eArchT-based engram inhibition groups (Figures 5I, 5S), which were normalized by their respective control group data. Engram cell reactivation in CA1 showed that memory retrieval is accompanied by enhanced cFos activation in several brain regions that are activated by natural memory recall (Figure 5I), which was a subset of the natural recall activity pattern. A similar observation was made for engram cell reactivation in BLA (Figure 5S). Importantly, when we statistically compared the brain-wide neuronal activity pattern following optogenetic engram cell reactivation in CA1 (Figure 5J) and BLA (Figure 5T) to the brain-wide neuronal activation pattern induced by natural recall, we observed a significant correlation between engram cell reactivation and natural recall for both brain region manipulations. These experiments demonstrate a comparable brain-wide activity pattern between natural memory recall and optogenetic engram cell reactivation, which suggests that optogenetic reactivation of engram cells recreates at least part of the pattern of brain-wide activity corresponding to natural memory recall. Further, engram cell inhibition in CA1 (Figure 5I) and BLA (Figure 5S) showed that memory retrieval impairments are accompanied by decreased cFos activation in several brain regions that are activated by natural memory recall, which is consistent with the observed behavioral disruptions. These data provide support for the idea that memory engram ensembles that are activated by learning are reactivated during recall of the specific memory (Semon, 1904; Tonegawa et al., 2015), and that this reactivation is a critical component of successful memory retrieval.

### Simultaneous chemogenetic reactivation of multiple engram cell ensembles

The brain-wide activity mapping using engram indices (Figures 1–3), memory recall induced by engram cell reactivation (Figure 4), and similar brain-wide neuronal activation pattern induced by natural memory recall and optogenetic activation-based recall (Figure 5), suggest that multiple engram cell ensembles distributed across the brain contribute to the efficient and specific memory retrieval process. Although the optogenetic reactivation of an individual engram cell ensemble results in significant levels of memory recall (Liu et al., 2012), the reactivation under naturalistic conditions would involve multiple engram cell ensembles. Indeed, optogenetic reactivation of a single engram cell ensemble does not reach the level of recall that is attained by natural recall (e.g., Liu et al., 2012). To investigate this issue, we took advantage of chemogenetic neuronal activation using the excitatory hM3Dq DREADDs receptor (Roth, 2016). This approach was selected since we could simultaneously activate multiple hM3Dq-expressing engram populations using the ligand clozapine-N-oxide (CNO). We developed a double virus approach expressing hM3Dq-mCherry in engram cells (Figure 6A). Our behavioral schedule included engram cell labeling during CFC training, followed by a natural memory recall test, and then finally CNO-induced engram cell reactivation in a neutral context on day 3 (Figure 6B). Since the first demonstration of engram cell reactivation-induced memory recall was performed in hippocampal DG (Liu et al., 2012), we initially attempted engram cell reactivation using the DREADDs system in hippocampal DG. Activating DG engram cells using CNO resulted in increased freezing behavior compared to mCherry control mice (Figure 6C), which validated this chemogenetic approach for engram cell reactivation.

**Figure 6.**
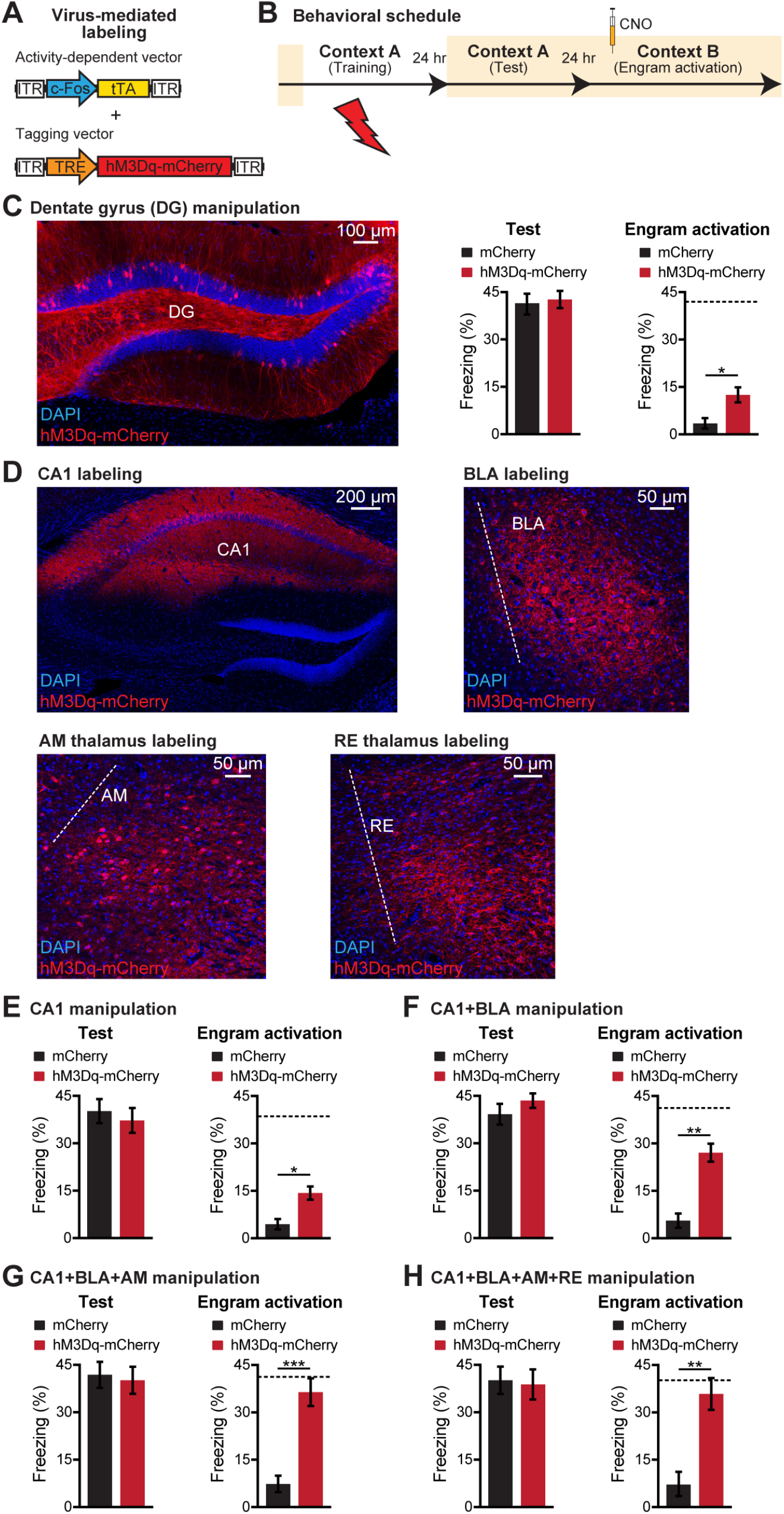
Simultaneous reactivation of multiple engram cell ensembles using DREADDs. (A) Virus-mediated cFos^+^ neuronal labeling strategy using a cocktail of c-Fos-tTA and TRE-hM3Dq-mCherry in wild-type mice. (B) Behavioral schedule. Beige shading signifies that mice were on DOX. Mice were taken off DOX 24 hrs before contextual fear conditioning (CFC). One day after training, a natural recall test was performed (Test). The next day, mice received CNO IP 45 min before a chemogenetic reactivation (Engram activation) session in a neutral context B. (C) Hippocampal DG section showing cFos^+^ neurons labeled with hM3Dq-mCherry using the two-virus approach (left). Natural memory recall (Test). Chemogenetic reactivation (Engram activation). mCherry (n = 10 mice) and hM3Dq-mCherry (n = 9 mice) groups. Dashed line indicates natural recall freezing level. (D) Hippocampal CA1, BLA, AM thalamus, and RE thalamus sections showing cFos^+^ neurons labeled with hM3Dq-mCherry using the two-virus approach. (E) Chemogenetic manipulation of CA1 engram cells. Natural recall (Test). Chemogenetic reactivation (Engram activation). mCherry (n = 8 mice) and hM3Dq-mCherry (n = 12 mice) groups. Dashed line indicates natural recall freezing level. (F) Chemogenetic manipulation of CA1 and BLA engram cells. Natural recall (Test). Chemogenetic reactivation (Engram activation). mCherry (n = 13 mice) and hM3Dq-mCherry (n = 11 mice) groups. Dashed line indicates natural recall freezing level. (G) Chemogenetic manipulation of CA1, BLA, and AM engram cells. Natural recall (Test). Chemogenetic reactivation (Engram activation). mCherry (n = 11 mice) and hM3Dq-mCherry (n = 11 mice) groups. Dashed line indicates natural recall freezing level. (H) Chemogenetic manipulation of CA1, BLA, AM, and RE engram cells. Natural recall (Test). Chemogenetic reactivation (Engram activation). mCherry (n = 10 mice) and hM3Dq-mCherry (n = 9 mice) groups. Dashed line indicates natural recall freezing level. Statistical comparisons are performed using unpaired *t* tests; *P < 0.05, **P < 0.01, ***P < 0.001. Data are presented as mean ± SEM.

We performed step-wise multiple engram cell ensemble reactivation experiments. Building on our findings using optogenetic engram cell reactivation (Figure 4), we reactivated a single engram cell ensemble (hippocampal CA1), two engram cell ensembles (CA1 and BLA) tagged in the same animals, three engram cell ensembles (CA1, BLA, and AM thalamus) tagged in the same animals, and four engram cell ensembles (CA1, BLA, AM and RE thalamus) tagged in the same animals (Figure 6D). Consistent with the corresponding optogenetic reactivation experiment (Figure 4F), CNO-induced CA1 engram cell reactivation resulted in memory recall (Figure 6E). Reactivation of two (Figure 6F) and three (Figure 6G) engram cell ensembles respectively conferred enhanced memory reactivation, as assessed by the percentage of time freezing. This finding indicates that the simultaneous reactivation of multiple engram cell ensembles results in greater memory reactivation than their subset(s). Interestingly, we did not observe further enhancement of memory reactivation by the simultaneous reactivation of four engram cell ensembles (Figure 6H), suggesting that there may be an upper limit in the capacity of memory engram cell reactivation to modulate behavioral outputs. Nevertheless, since the simultaneous reactivation of three and four engram cell ensembles resulted in memory recall strength that was comparable to natural memory recall, this may reflect a mechanism by which the brain adjusts the strength of memory recall depending on the importance of the memory for current and future decisions.

## DISCUSSION

### Previous attempts of brain-wide engram ensemble mapping

Generating a brain-wide map of anatomical regions crucially responsible for high level cognitive functions and resulting behaviors, such as memory acquisition and expression, is technically demanding. Even if a method is available, it is usually tedious and impractical to apply to all brain regions. Therefore, previous studies relied on an indirect method and determined the brain-wide distribution of neuronal ensembles activated by a variety of behavioral paradigms and supplemented the resulting activity map with additional data analysis and/or experiments. For instance, focusing on remote fear memory retrieval, one study employed immediate early gene (cFos) activation in tissue sections to quantify the activity of 84 brain regions (Vetere et al., 2017). This study further showed that co-active regions during remote memory retrieval are more important for network function using chemogenetic neuronal inhibition and computational modeling. However, laborious sample processing and information loss during manual tissue sectioning limited the applicability of such approaches. In three other studies, the advent of tissue clearing techniques enabled holistic visualization of activated ensembles in intact mouse brain samples (Chung et al., 2013; Renier et al., 2014). The first study combined iDISCO+ clearing with brain-wide cFos staining to investigate the effects of haloperidol administration, whisker stimulation, and parental behavior (Renier et al., 2016). They identified new cortical regions activated by whisker stimulation during exploration and uncovered differentially activated regions during parental behavior. However, this study did not perform functional manipulations of the activated cell ensembles and therefore it is not certain that they hold engrams. The second study used the CLARITY method with genetically-labeled Arc-TRAP mice to investigate the effects of foot shocks and cocaine administration (Ye et al., 2016). The analysis was restricted to 9 brain regions in the forebrain, and the authors focused on prefrontal cortex ensembles using optogenetic manipulations. This study showed for the first time that different cell populations in the prefrontal cortex represent negative- and positive-valence experiences. A third study used iDISCO+ clearing with genetically-labeled Fos-TRAP2 mice to investigate neurons activated by fear conditioning in 200 regions, with a focus on remote memory retrieval (DeNardo et al., 2019). Although functional manipulations were limited to prefrontal cortex ensembles, this study found that some cFos^+^ neurons were indeed engram cells.

### A three-step approach for improved brain-wide mapping of engram cells of a specific memory

In the present study, we performed brain-wide high-throughput screening of putative engram ensembles of a contextual fear memory by combining SHIELD-based brain phenotyping (Park et al., 2018) and engram index analyses. Preservation of both protein fluorescence and tissue architecture by SHIELD enabled automated 3D analysis of fluorescent protein-tagged mouse brains and the generation of brain-wide activity maps of 409 regions.

The engram index is based on the concept that engrams are held by neuronal ensembles that are activated by learning and are reactivated to support recall (Josselyn et al., 2015; Semon, 1904; Tonegawa et al., 2015). In other words, a brain region containing engram cells for a specific memory will show high levels of activation during both memory encoding and recall, whereas this region will show low levels of activation in home cage or a neutral context. To reflect these criteria, the engram index equation has two components. The first component (i.e., numerator of the equation) computes the increase in average number of cFos-activated neurons for a given brain region during memory encoding as compared to neutral context exposure. By penalizing brain regions with high activation levels in the neutral context, this index can be used to rank brain regions that specifically contribute to context-fear associations and not those that only encode contextual representations. The second component (i.e., denominator of the equation) computes the difference between activation levels during memory encoding and recall. In doing so, this component decreases the engram index for brain regions with high activation levels during either encoding or recall, which also helps exclude regions that are primarily responsive to the foot shocks themselves (i.e., false positive candidate engram regions). To summarize, brain regions with high indices indicate that these structures satisfy two criteria: (a) a clear increase in activation levels specifically during context-fear associations as compared to context only experiences, and (b) high similarity between encoding and recall activation levels, which is in agreement with the concept that engram cells are activated during learning and reactivated during recall. Most known brain regions that have previously been demonstrated to hold engrams by optogenetic and other manipulations (e.g., hippocampal dentate gyrus, CA3, and basolateral amygdala) showed high engram index values, supporting our hypothesis that a brain region with a high index is likely to hold an engram cell ensemble. One limitation of this index, however, is that it cannot be used to identify silent engram cells, which are formed in certain brain regions during encoding but are not reactivated by natural recall cues for recall although they can be optogenetically reactivated (Roy et al., 2017a; Ryan et al., 2015; Tonegawa et al., 2018). A second limitation of this index is that it is limited to brain regions in which cFos is expressed, which clearly is the case for the majority of regions. The brain regions that gave high engram indices are likely to carry engrams and thereby were selected for the cumbersome but powerful functional intervention experiments to confirm the existence of engrams. This permitted us to identify two new brain regions as holders of engram cell ensembles, anteromedial thalamus (AM) and nucleus reuniens thalamus (RE).

### Potential role of thalamic engram cell ensembles

Since AM thalamus receives projections from the medial hypothalamic circuit responsible for threat processing, and has reciprocal connectivity with multiple cortical regions, it is likely that AM thalamus engram cells convey cortico-hypothalamic information to hippocampal circuitry for the generation of context-specific, high-valence memories (de Lima et al., 2017). On the other hand, RE thalamus has been reported to play a crucial role in hippocampal-dependent encoding of spatial memories, particularly contributing to the discrimination of similar environments (Ramanathan et al., 2018). Therefore, RE engram cells may enhance the specificity of contextual memory retrieval by regulating the online discrimination ability of the animal. It would be worthwhile to further identify engram cell ensembles in other thalamic nuclei and determine their unique roles in memory processes.

### Memory storage in distributed engram cell ensembles

The concept that a given memory is stored not just in a single engram cell ensemble but in learning-induced enduring changes in functionally connected multiple neuronal ensembles was predicted by Richard Semon (“engram complex”) (Semon, 1904) and Donald Hebb (“neurons that fire together wire together”) (Hebb, 1949). The experimental evidence for this concept came from an analysis of engram cells from multiple hippocampal subfields and the amygdala (Choi et al., 2018; Redondo et al., 2014; Ryan et al., 2015) and has since been supported by activity mapping studies (DeNardo et al., 2019; Renier et al., 2016; Vetere et al., 2017; Ye et al., 2016).

The present study provides two additional types of support for the engram complex hypothesis. First, we have shown that optogenetic activation of a single engram cell ensemble (the one in CA1 or BLA) can activate a set of other engram ensembles or candidate engram ensembles, demonstrating functional connectivity of engram ensembles across multiple brain regions. Second, the Semon hypothesis assumed that engrams scattered in multiple brain regions will be reactivated simultaneously by natural recall cues. In contrast, optogenetic memory retrieval is usually accomplished by reactivation of a single engram ensemble but the resulting level of recall (for instance, level of freezing during recall of a contextual fear memory) never reaches the level attained by natural recall cues. In the present study, we showed that recall level correlated with the number of engram ensembles reactivated simultaneously by delivery of the chemogenetic ligand, supporting the hypothesis that memory recall is accomplished by simultaneous reactivation of multiple engram ensembles across brain regions.

The distributed nature of engram cell ensembles for a specific memory has led to the suggestion that the memory engram within an individual brain region may contribute a subset of the overall memory information (Eichenbaum, 2016; Roy et al., 2017b). For example, hippocampal engrams are thought to primarily contribute contextual information by acting as an index for cortical memories of various sensory modalities (Teyler and DiScenna, 1986; Teyler and Rudy, 2007), whereas amygdala engrams hold valence information for a given experience (Janak and Tye, 2015; Morrison and Salzman, 2010; Redondo et al., 2014; Tovote et al., 2015). In addition, cortical engrams such as those in the retrosplenial and prefrontal cortices may support spatial navigation (Vann et al., 2009) and top-down control of memory retrieval (Tomita et al., 1999), respectively. Further, engrams in auditory (Fritz et al., 2005) and olfactory cortices (Shakhawat et al., 2014) may support auditory recognition memory and odor-induced learned behaviors, respectively.

In conclusion, this study provides evidence supporting the concept that a given memory is stored in functionally connected engram ensembles distributed broadly in multiple brain regions. Despite some caveats, our three-step strategy has provided to-date the most comprehensive mapping of engrams- and high probability engram-holding brain regions. Future studies can take advantage of this resource to generate a more extensive map of engram cell ensembles including the identification of their functional connectivity as well as the mnemonic functions of individual ensembles.

## AUTHOR CONTRIBUTIONS

D.S.R., Y.-G.P., S.K.O., K.C., and S.T. contributed to the study design. D.S.R., Y.-G.P., S.K.O., J.H.C., H.C., and J.M. contributed to the data collection. D.S.R., Y.-G.P., S.K.O., and S.T. contributed to the data interpretation. D.S.R., S.K.O., and J.M. conducted the surgeries, behavior experiments, and histological analyses. Y.-G.P., J.H.C., H.C., and L.K. conducted the tissue clearing, light-sheet microscopy, and computational analyses. D.S.R., Y.-G.P., S.K.O., K.C., and S.T. wrote the paper. All authors discussed and commented on the manuscript.

## ACKNOWLEDGEMENTS

We thank S. Huang, L. Smith, F. Bushard, A. Hamalian, D. King, C. Ragion, W. Yu, and C. Lovett for help with experiments; C. Sun, S. Muralidhar, and Q. Ferry for comments; A. Schroeder for proofreading; and all members of the Chung and Tonegawa laboratories for their support. This work was supported by the Burroughs Wellcome Fund Career Award at the Scientific Interface, Searle Scholars Program, Packard Award in Science and Engineering, NARSAD Young Investigator Award, McKnight Foundation Technology Award, JPB Foundation (PIIF and PNDRF), NCSOFT Cultural Foundation, the Institute for Basic Science IBS-R026-D1, and NIH (1-DP2-ES027992) (to K.C.), and by the RIKEN Center for Brain Science, Howard Hughes Medical Institute, and JPB Foundation (to S.T.).

