## Supplementary Materials for "Brain-wide mapping of contextual fear memory engram ensembles supports the dispersed engram complex hypothesis"

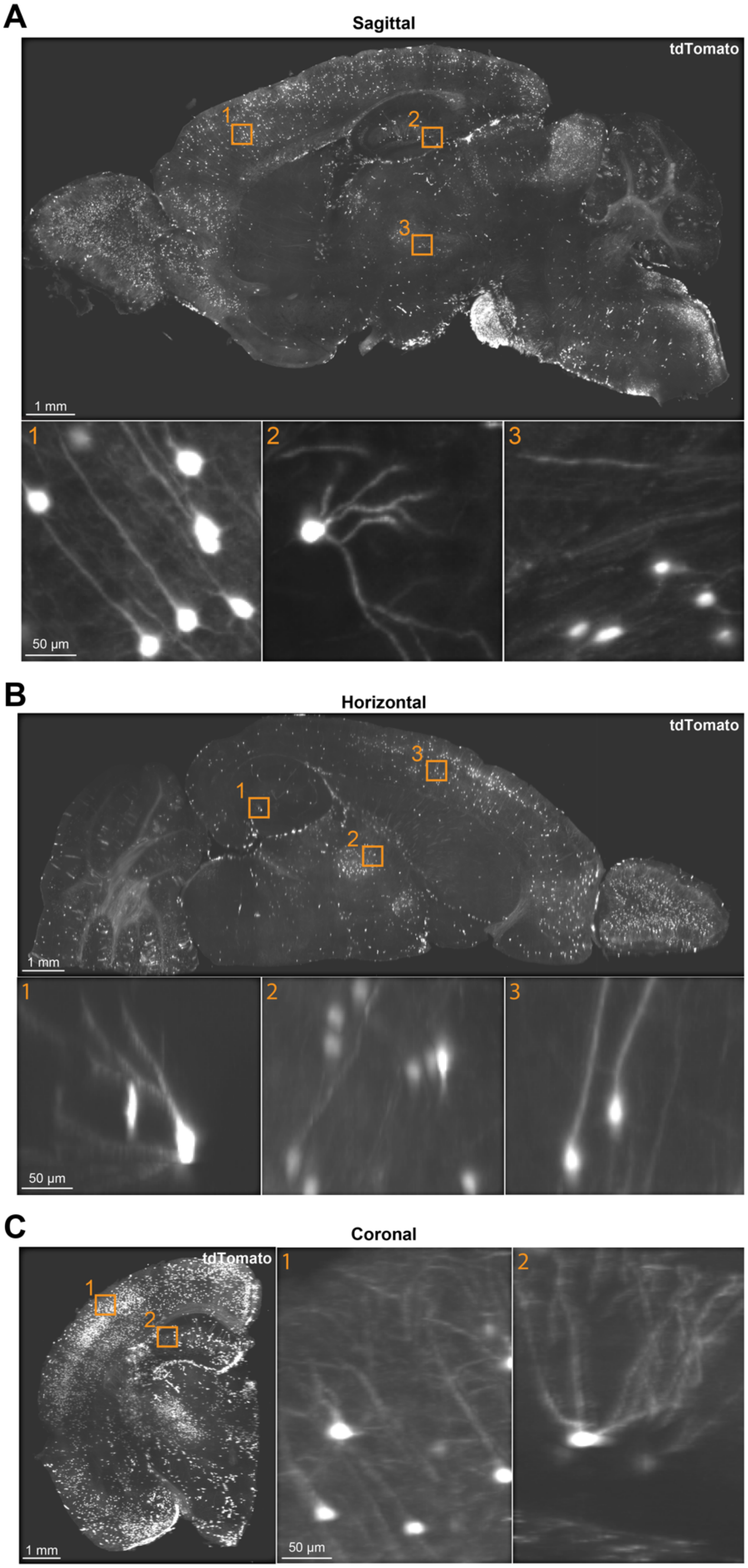

**Figure S1. Orthogonal views of a CFC brain, Related to Figure 1**

(A-C) 3D reconstructions were used to quantify brain-wide $cFos^+$ neuronal ensembles labeled during different behavioral epochs. Using a contextual fear conditioning (CFC) brain as an example, we extracted 2D orthogonal views from these 3D reconstructions. Sagittal views (A), horizontal views (B), and coronal views (C) demonstrate the single cell resolution achieved using our brain-wide activity mapping approach.

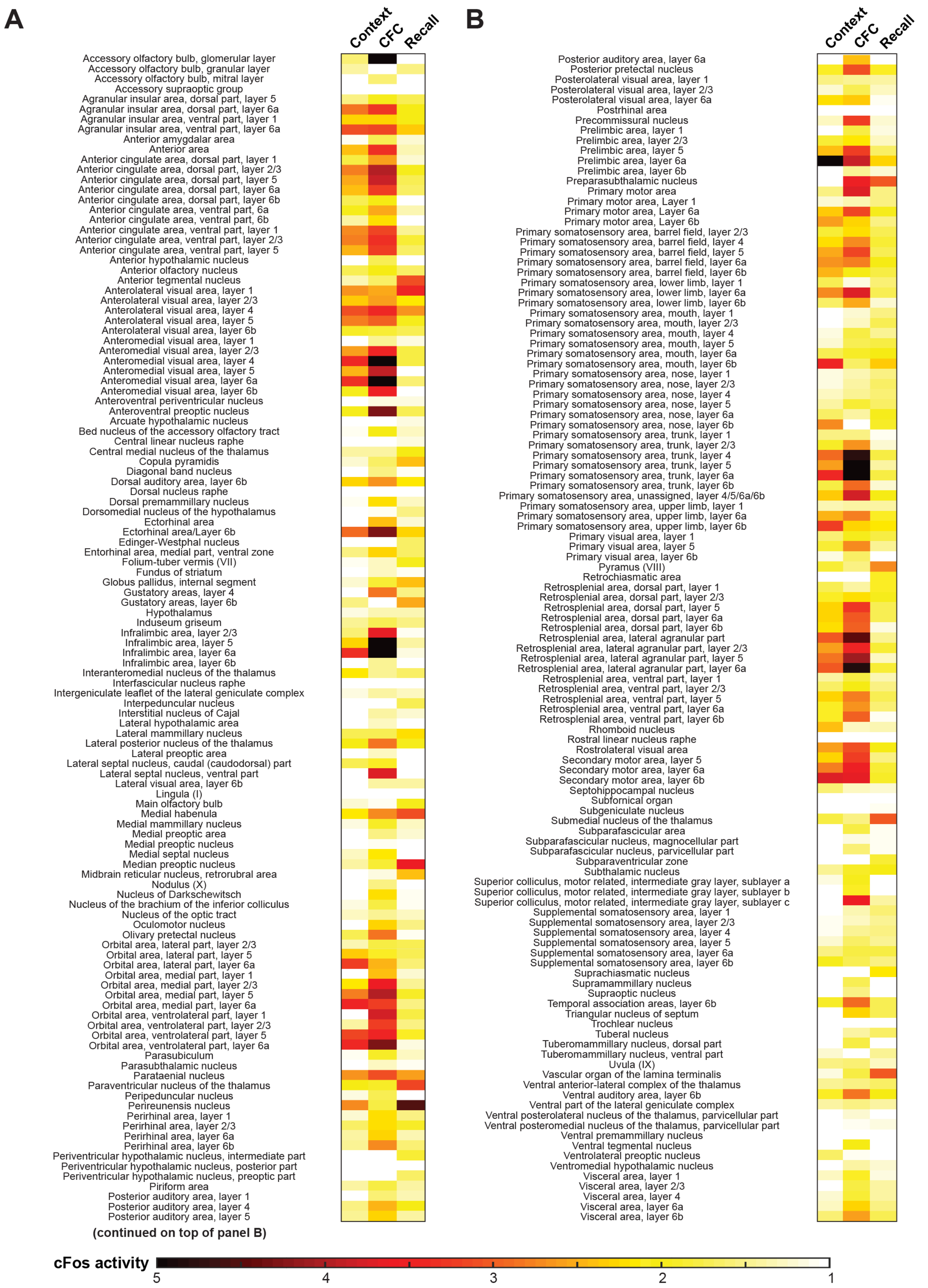
A
Context
CFC
Recall
Accessory olfactory bulb, glomerular layer
Accessory olfactory bulb, granular layer
Accessory olfactory bulb, mitral layer
Accessory supraoptic group
Agranular insular area, dorsal part, layer 5
Agranular insular area, dorsal part, layer 6a
Agranular insular area, ventral part, layer 1
Agranular insular area, ventral part, layer 6a
Anterior amygdalar area
Anterior area
Anterior cingulate area, dorsal part, layer 1
Anterior cingulate area, dorsal part, layer 2/3
Anterior cingulate area, dorsal part, layer 5
Anterior cingulate area, dorsal part, layer 6a
Anterior cingulate area, dorsal part, layer 6b
Anterior cingulate area, ventral part, 6a
Anterior cingulate area, ventral part, 6b
Anterior cingulate area, ventral part, layer 1
Anterior cingulate area, ventral part, layer 2/3
Anterior cingulate area, ventral part, layer 5
Anterior hypothalamic nucleus
Anterior olfactory nucleus
Anterior tegmental nucleus
Anterolateral visual area, layer 1
Anterolateral visual area, layer 2/3
Anterolateral visual area, layer 4
Anterolateral visual area, layer 5
Anterolateral visual area, layer 6b
Anteromedial visual area, layer 1
Anteromedial visual area, layer 2/3
Anteromedial visual area, layer 4
Anteromedial visual area, layer 5
Anteromedial visual area, layer 6a
Anteromedial visual area, layer 6b
Anteroventral periventricular nucleus
Anteroventral preoptic nucleus
Arcuate hypothalamic nucleus
Bed nucleus of the accessory olfactory tract
Central linear nucleus raphe
Central medial nucleus of the thalamus
Copula pyramidis
Diagonal band nucleus
Dorsal auditory area, layer 6b
Dorsal nucleus raphe
Dorsal premammillary nucleus
Dorsomedial nucleus of the hypothalamus
Ectorhinal area
Ectorhinal area/Layer 6b
Edinger-Westphal nucleus
Entorhinal area, medial part, ventral zone
Folium-tuber vermis (VII)
Fundus of striatum
Globus pallidus, internal segment
Gustatory areas, layer 4
Gustatory areas, layer 6b
Hypothalamus
Induseum griseum
Infralimbic area, layer 2/3
Infralimbic area, layer 5
Infralimbic area, layer 6a
Infralimbic area, layer 6b
Interanteromedial nucleus of the thalamus
Interfascicular nucleus raphe
Intergeniculate leaflet of the lateral geniculate complex
Interpeduncular nucleus
Interstitial nucleus of Cajal
Lateral hypothalamic area
Lateral mammillary nucleus
Lateral posterior nucleus of the thalamus
Lateral preoptic area
Lateral septal nucleus, caudal (caudodorsal) part
Lateral septal nucleus, ventral part
Lateral visual area, layer 6b
Lingula (I)
Main olfactory bulb
Medial habenula
Medial mammillary nucleus
Medial preoptic area
Medial preoptic nucleus
Medial septal nucleus
Median preoptic nucleus
Midbrain reticular nucleus, retrorubral area
Nodulus (X)
Nucleus of Darkschewitsch
Nucleus of the brachium of the inferior colliculus
Nucleus of the optic tract
Oculomotor nucleus
Olivary pretectal nucleus
Orbital area, lateral part, layer 2/3
Orbital area, lateral part, layer 5
Orbital area, lateral part, layer 6a
Orbital area, medial part, layer 1
Orbital area, medial part, layer 2/3
Orbital area, medial part, layer 5
Orbital area, medial part, layer 6a
Orbital area, ventrolateral part, layer 1
Orbital area, ventrolateral part, layer 2/3
Orbital area, ventrolateral part, layer 5
Orbital area, ventrolateral part, layer 6a
Parasubiculum
Parasubthalamic nucleus
Parataenial nucleus
Paraventricular nucleus of the thalamus
Peripeduncular nucleus
Perireunensis nucleus
Perirhinal area, layer 1
Perirhinal area, layer 2/3
Perirhinal area, layer 6a
Perirhinal area, layer 6b
Periventricular hypothalamic nucleus, intermediate part
Periventricular hypothalamic nucleus, posterior part
Periventricular hypothalamic nucleus, preoptic part
Piriform area
Posterior auditory area, layer 1
Posterior auditory area, layer 4
Posterior auditory area, layer 5
(continued on top of panel B)
B
Context
CFC
Recall
Posterior auditory area, layer 6a
Posterior pretectal nucleus
Posterolateral visual area, layer 1
Posterolateral visual area, layer 2/3
Posterolateral visual area, layer 6a
Postrhinal area
Precommissural nucleus
Prelimbic area, layer 1
Prelimbic area, layer 2/3
Prelimbic area, layer 5
Prelimbic area, layer 6a
Prelimbic area, layer 6b
Preparasubthalamic nucleus
Primary motor area
Primary motor area, Layer 1
Primary motor area, Layer 6a
Primary motor area, Layer 6b
Primary somatosensory area, barrel field, layer 2/3
Primary somatosensory area, barrel field, layer 4
Primary somatosensory area, barrel field, layer 5
Primary somatosensory area, barrel field, layer 6a
Primary somatosensory area, barrel field, layer 6b
Primary somatosensory area, lower limb, layer 1
Primary somatosensory area, lower limb, layer 6a
Primary somatosensory area, lower limb, layer 6b
Primary somatosensory area, mouth, layer 1
Primary somatosensory area, mouth, layer 2/3
Primary somatosensory area, mouth, layer 4
Primary somatosensory area, mouth, layer 5
Primary somatosensory area, mouth, layer 6a
Primary somatosensory area, mouth, layer 6b
Primary somatosensory area, nose, layer 1
Primary somatosensory area, nose, layer 2/3
Primary somatosensory area, nose, layer 4
Primary somatosensory area, nose, layer 5
Primary somatosensory area, nose, layer 6a
Primary somatosensory area, nose, layer 6b
Primary somatosensory area, trunk, layer 1
Primary somatosensory area, trunk, layer 2/3
Primary somatosensory area, trunk, layer 4
Primary somatosensory area, trunk, layer 5
Primary somatosensory area, trunk, layer 6a
Primary somatosensory area, trunk, layer 6b
Primary somatosensory area, unassigned, layer 4/5/6a/6b
Primary somatosensory area, upper limb, layer 1
Primary somatosensory area, upper limb, layer 6a
Primary somatosensory area, upper limb, layer 6b
Primary visual area, layer 1
Primary visual area, layer 5
Primary visual area, layer 6b
Pyramus (VIII)
Retrochiasmatic area
Retrosplenial area, dorsal part, layer 1
Retrosplenial area, dorsal part, layer 2/3
Retrosplenial area, dorsal part, layer 5
Retrosplenial area, dorsal part, layer 6a
Retrosplenial area, dorsal part, layer 6b
Retrosplenial area, lateral agranular part
Retrosplenial area, lateral agranular part, layer 2/3
Retrosplenial area, lateral agranular part, layer 5
Retrosplenial area, lateral agranular part, layer 6a
Retrosplenial area, ventral part, layer 1
Retrosplenial area, ventral part, layer 2/3
Retrosplenial area, ventral part, layer 5
Retrosplenial area, ventral part, layer 6a
Retrosplenial area, ventral part, layer 6b
Rhomboid nucleus
Rostral linear nucleus raphe
Rostrolateral visual area
Secondary motor area, layer 5
Secondary motor area, layer 6a
Secondary motor area, layer 6b
Septohippocampal nucleus
Subfornical organ
Subgeniculate nucleus
Submedial nucleus of the thalamus
Subparafascicular area
Subparafascicular nucleus, magnocellular part
Subparafascicular nucleus, parvicellular part
Subparaventricular zone
Subthalamic nucleus
Superior colliculus, motor related, intermediate gray layer, sublayer a
Superior colliculus, motor related, intermediate gray layer, sublayer b
Superior colliculus, motor related, intermediate gray layer, sublayer c
Supplemental somatosensory area, layer 1
Supplemental somatosensory area, layer 2/3
Supplemental somatosensory area, layer 4
Supplemental somatosensory area, layer 5
Supplemental somatosensory area, layer 6a
Supplemental somatosensory area, layer 6b
Suprachiasmatic nucleus
Supramammillary nucleus
Supraoptic nucleus
Temporal association areas, layer 6b
Triangular nucleus of septum
Trochlear nucleus
Tuberal nucleus
Tuberomammillary nucleus, dorsal part
Tuberomammillary nucleus, ventral part
Uvula (IX)
Vascular organ of the lamina terminalis
Ventral anterior-lateral complex of the thalamus
Ventral auditory area, layer 6b
Ventral part of the lateral geniculate complex
Ventral posterolateral nucleus of the thalamus, parvicellular part
Ventral posteromedial nucleus of the thalamus, parvicellular part
Ventral premammillary nucleus
Ventral tegmental nucleus
Ventrolateral preoptic nucleus
Ventromedial hypothalamic nucleus
Visceral area, layer 1
Visceral area, layer 2/3
Visceral area, layer 4
Visceral area, layer 6a
Visceral area, layer 6b
cFos activity
5
4
3
2
1

**Figure S2. Activity mapping results for brain structures with non-significant engram indices, Related to Figure 2**

(A and B) Figure 2 shows a rank-ordered list of brain-wide engram indices. The neuronal activity across behavioral groups is shown in this supplemental figure for 231 brain regions and their subdivisions that are not included in Figure 2 due to their non-significant indices (n = 4 mice each for home cage and CFC groups, n = 3 mice each for context and recall groups). Color-coded scale bar represents fold increases in the numbers of activated neurons relative to home cage.

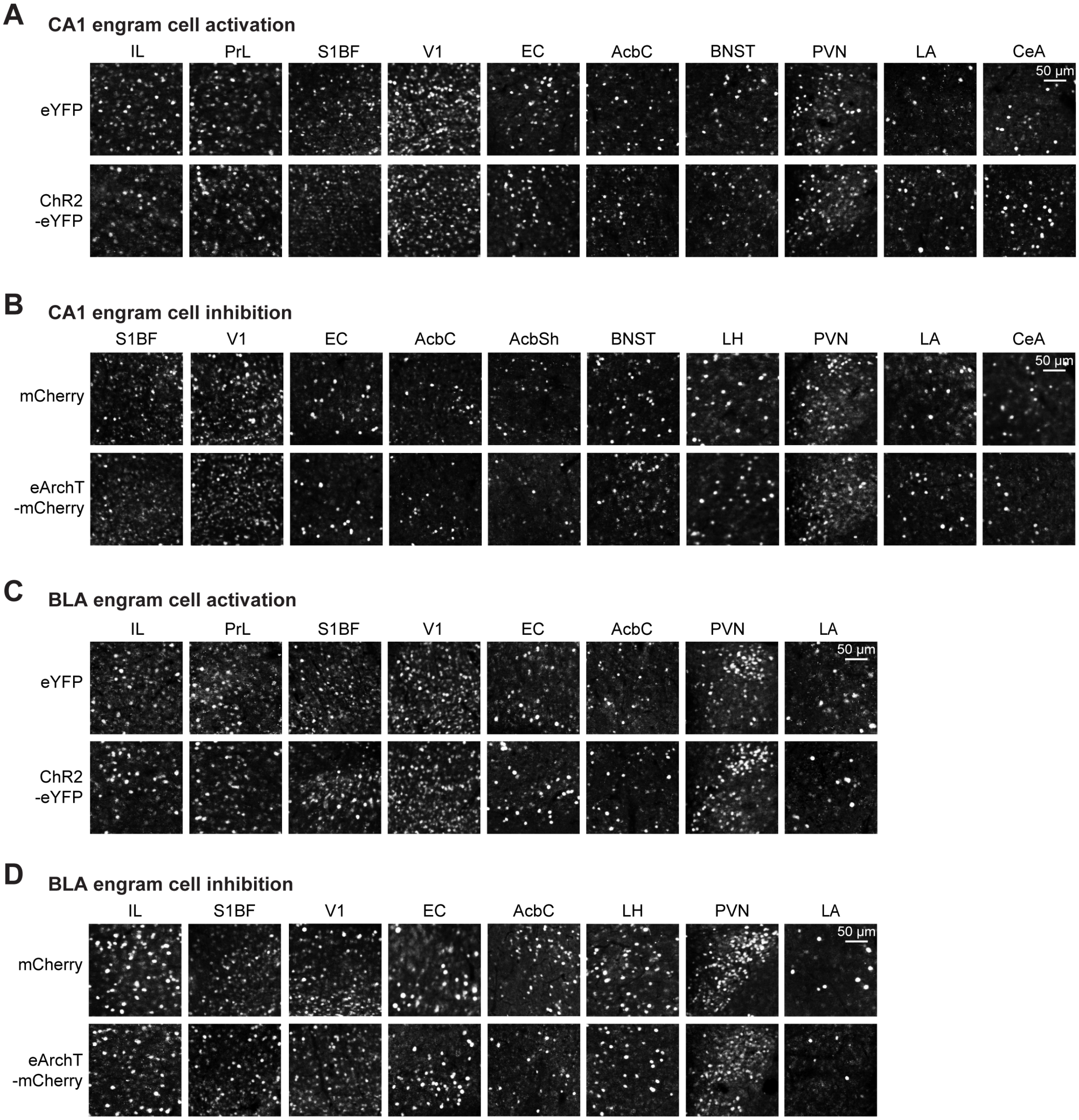


**Figure S3. cFos$^+$ neurons across brain regions following CA1 or BLA engram cell manipulations, Related to Figure 5**

(A and B) cFos$^+$ neurons in representative brain regions from CA1 engram cell activation (A) or CA1 engram cell inhibition (B) mice. eYFP and mCherry groups (top row), ChR2-eYFP and eArchT-mCherry groups (bottom row). See Figure 5I.

(C and D) cFos$^+$ neurons in representative brain regions from BLA engram cell activation (C) or BLA engram cell inhibition (D) mice. eYFP and mCherry groups (top row), ChR2-eYFP and eArchT-mCherry groups (bottom row). See Figure 5S.

**Movie S1. Brain-wide labeling of cFos$^+$ neuronal ensembles in home cage and CFC mice, Related to Figures 1-3**

Representative movie of Fos-TRAP mice crossed with Cre-dependent tdTomato reporter mice, which was used to label cFos$^+$ neurons in the home cage condition (Home cage) or the contextual fear training condition (CFC). Movie is shown in sagittal view.

**Movie S2. cFos$^+$ neuronal ensembles in the hippocampus, Related to Figures 1-3**

Representative high-resolution movie of a Fos-TRAP mouse crossed with a Cre-dependent tdTomato reporter mouse, which shows cFos$^+$ neurons in an optical section focused on the hippocampus of a contextual fear memory recall-labeled animal. Movie is shown in sagittal view.

**Movie S3. Color-coded brain-wide engram indices, Related to Figure 2**

Representative movie of $cFos^+$ neuronal ensembles in each brain region color-coded for their respective engram index values (ranging from 0.03 to 3.16). Lower engram indices are closer to the blue color, whereas higher engram indices are closer to the red color. Movie is shown in sagittal view.

#### Table S1. List of 409 brain regions analyzed in the brain-wide activity maps, Related to Figures 1 and 2

Accessory olfactory bulb, granular layer
Accessory olfactory bulb, mitral layer
Accessory supraoptic group
Agranular insular area, dorsal part, layer 1
Agranular insular area, dorsal part, layer 2/3
Agranular insular area, dorsal part, layer 5
Agranular insular area, dorsal part, layer 6a
Agranular insular area, posterior part, layer 1
Agranular insular area, posterior part, layer 2/3
Agranular insular area, posterior part, layer 5
Agranular insular area, posterior part, layer 6a
Agranular insular area, ventral part, layer 1
Agranular insular area, ventral part, layer 2/3
Agranular insular area, ventral part, layer 5
Agranular insular area, ventral part, layer 6a
Ansiform lobule
Anterior amygdalar area
Anterior area
Anterior cingulate area, dorsal part, layer 1
Anterior cingulate area, dorsal part, layer 2/3
Anterior cingulate area, dorsal part, layer 5
Anterior cingulate area, dorsal part, layer 6a
Anterior cingulate area, dorsal part, layer 6b
Anterior cingulate area, ventral part, 6a
Anterior cingulate area, ventral part, 6b
Anterior cingulate area, ventral part, layer 1
Anterior cingulate area, ventral part, layer 2/3
Anterior cingulate area, ventral part, layer 5
Anterior hypothalamic nucleus
Anterior olfactory nucleus
Anterior pretectal nucleus
Anterior tegmental nucleus
Anterodorsal nucleus
Anterodorsal preoptic nucleus
Anterolateral visual area, layer 1
Anterolateral visual area, layer 2/3
Anterolateral visual area, layer 4
Anterolateral visual area, layer 5
Anterolateral visual area, layer 6a
Anterolateral visual area, layer 6b
Anteromedial nucleus
Anteromedial visual area, layer 1
Anteromedial visual area, layer 2/3
Anteromedial visual area, layer 4
Anteromedial visual area, layer 5
Anteromedial visual area, layer 6a
Anteromedial visual area, layer 6b
Anteroventral nucleus of thalamus
Anteroventral periventricular nucleus
Anteroventral preoptic nucleus
Arcuate hypothalamic nucleus
Basolateral amygdalar nucleus, anterior part
Basolateral amygdalar nucleus, posterior part
Basolateral amygdalar nucleus, ventral part
Basomedial amygdalar nucleus
Bed nuclei of the stria terminalis
Bed nucleus of the accessory olfactory tract
Caudoputamen
Central amygdalar nucleus
Central lateral nucleus of the thalamus
Central linear nucleus raphe
Central lobule
Central medial nucleus of the thalamus
Claustrum
Copula pyramidis
Cortical amygdalar area, anterior part
Cortical amygdalar area, posterior part
Cortical subplate
Culmen
Cuneiform nucleus
Declive (VI)
Dentate gyrus, granule cell layer
Dentate gyrus, molecular layer
Dentate gyrus, polymorph layer
Dentate nucleus
Diagonal band nucleus
Dorsal auditory area, layer 1
Dorsal auditory area, layer 2/3
Dorsal auditory area, layer 4
Dorsal auditory area, layer 5
Dorsal auditory area, layer 6a
Dorsal auditory area, layer 6b
Dorsal nucleus raphe
Dorsal part of the lateral geniculate complex
Dorsal peduncular area
Dorsal premammillary nucleus
Dorsomedial nucleus of the hypothalamus
Ectorhinal area
Ectorhinal area/Layer 1
Ectorhinal area/Layer 2/3
Ectorhinal area/Layer 5
Ectorhinal area/Layer 6a
Ectorhinal area/Layer 6b
Edinger-Westphal nucleus
Endopiriform nucleus, dorsal part
Endopiriform nucleus, ventral part
Entorhinal area, lateral part
Entorhinal area, medial part, dorsal zone
Entorhinal area, medial part, ventral zone
Fasciola cinerea
Fastigial nucleus
Field CA1
Field CA2
Field CA3
Fields of Forel
Flocculus
Folium-tuber vermis (VII)
Frontal pole, layer 1
Frontal pole, layer 2/3
Fundus of striatum
Globus pallidus, external segment
Globus pallidus, internal segment
Gustatory areas, layer 1
Gustatory areas, layer 2/3
Gustatory areas, layer 4
Gustatory areas, layer 5
Gustatory areas, layer 6a
Gustatory areas, layer 6b
Hippocampal formation
Hypothalamus
Induseum griseum
Inferior colliculus
Infralimbic area, layer 1
Infralimbic area, layer 2/3
Infralimbic area, layer 5
Infralimbic area, layer 6a
Infralimbic area, layer 6b
Interanterodorsal nucleus of the thalamus
Interanteromedial nucleus of the thalamus
Intercalated amygdalar nucleus
Interfascicular nucleus raphe
Intergeniculate leaflet of the lateral geniculate complex
Intermediodorsal nucleus of the thalamus
Interpeduncular nucleus
Interposed nucleus
Interstitial nucleus of Cajal

#### Table S1 (continued)

| | |
|---|---|
| Lateral amygdalar nucleus | Parataenial nucleus |
| Lateral dorsal nucleus of thalamus | Paraventricular hypothalamic nucleus |
| Lateral habenula | Paraventricular hypothalamic nucleus, descending division |
| Lateral hypothalamic area | Paraventricular nucleus of the thalamus |
| Lateral mammillary nucleus | Pedunculopontine nucleus |
| Lateral posterior nucleus of the thalamus | Periaqueductal gray |
| Lateral preoptic area | Peripeduncular nucleus |
| Lateral septal nucleus, caudal (caudodorsal) part | Perireunensis nucleus |
| Lateral septal nucleus, rostral (rostroventral) part | Perirhinal area, layer 1 |
| Lateral septal nucleus, ventral part | Perirhinal area, layer 2/3 |
| Lateral visual area, layer 1 | Perirhinal area, layer 5 |
| Lateral visual area, layer 2/3 | Perirhinal area, layer 6a |
| Lateral visual area, layer 4 | Perirhinal area, layer 6b |
| Lateral visual area, layer 5 | Periventricular hypothalamic nucleus, intermediate part |
| Lateral visual area, layer 6a | Periventricular hypothalamic nucleus, posterior part |
| Lateral visual area, layer 6b | Periventricular hypothalamic nucleus, preoptic part |
| Laterointermediate area | Piriform area |
| Lingula (I) | Piriform-amygdalar area |
| Magnocellular nucleus | Posterior amygdalar nucleus |
| Main olfactory bulb | Posterior auditory area, layer 1 |
| Medial amygdalar nucleus | Posterior auditory area, layer 2/3 |
| Medial geniculate complex | Posterior auditory area, layer 4 |
| Medial habenula | Posterior auditory area, layer 5 |
| Medial mammillary nucleus | Posterior auditory area, layer 6a |
| Medial preoptic area | Posterior complex of the thalamus |
| Medial preoptic nucleus | Posterior hypothalamic nucleus |
| Medial pretectal area | Posterior limiting nucleus of the thalamus |
| Medial septal nucleus | Posterior pretectal nucleus |
| Median preoptic nucleus | Posterolateral visual area, layer 1 |
| Mediodorsal nucleus of thalamus | Posterolateral visual area, layer 2/3 |
| Midbrain | Posterolateral visual area, layer 4 |
| Midbrain reticular nucleus | Posterolateral visual area, layer 5 |
| Midbrain reticular nucleus, retrorubral area | Posterolateral visual area, layer 6a |
| Midbrain trigeminal nucleus | Postpiriform transition area |
| Nodulus (X) | Postrhinal area |
| Nucleus accumbens | Postsubiculum |
| Nucleus of Darkschewitsch | Precommissural nucleus |
| Nucleus of reunions | Prelimbic area, layer 1 |
| Nucleus of the brachium of the inferior colliculus | Prelimbic area, layer 2/3 |
| Nucleus of the lateral olfactory tract, layer 3 | Prelimbic area, layer 5 |
| Nucleus of the lateral olfactory tract, molecular layer | Prelimbic area, layer 6a |
| Nucleus of the optic tract | Prelimbic area, layer 6b |
| Nucleus of the posterior commissure | Preparasubthalamic nucleus |
| Nucleus sagulum | Presubiculum |
| Oculomotor nucleus | Primary auditory area, layer 1 |
| Olfactory areas | Primary auditory area, layer 2/3 |
| Olfactory tubercle | Primary auditory area, layer 4 |
| Olivary pretectal nucleus | Primary auditory area, layer 5 |
| Orbital area, lateral part, layer 1 | Primary auditory area, layer 6a |
| Orbital area, lateral part, layer 2/3 | Primary auditory area, layer 6b |
| Orbital area, lateral part, layer 5 | Primary motor area |
| Orbital area, lateral part, layer 6a | Primary motor area, Layer 1 |
| Orbital area, medial part, layer 1 | Primary motor area, Layer 2/3 |
| Orbital area, medial part, layer 2/3 | Primary motor area, Layer 5 |
| Orbital area, medial part, layer 5 | Primary motor area, Layer 6a |
| Orbital area, medial part, layer 6a | Primary motor area, Layer 6b |
| Orbital area, ventrolateral part, layer 1 | Primary somatosensory area, barrel field, layer 1 |
| Orbital area, ventrolateral part, layer 2/3 | Primary somatosensory area, barrel field, layer 2/3 |
| Orbital area, ventrolateral part, layer 5 | Primary somatosensory area, barrel field, layer 4 |
| Orbital area, ventrolateral part, layer 6a | Primary somatosensory area, barrel field, layer 5 |
| Pallidum | Primary somatosensory area, barrel field, layer 6a |
| Parabigeminal nucleus | Primary somatosensory area, barrel field, layer 6b |
| Paracentral nucleus | Primary somatosensory area, lower limb, layer 1 |
| Parafascicular nucleus | Primary somatosensory area, lower limb, layer 2/3 |
| Paraflocculus | Primary somatosensory area, lower limb, layer 4 |
| Paramedian lobule | Primary somatosensory area, lower limb, layer 5 |
| Parasubiculum | Primary somatosensory area, lower limb, layer 6a |
| Parasubthalamic nucleus | Primary somatosensory area, lower limb, layer 6b |

#### Table S1 (continued)

| | |
|---|---|
| Primary somatosensory area, mouth, layer 1 | Submedial nucleus of the thalamus |
| Primary somatosensory area, mouth, layer 2/3 | Subparafascicular area |
| Primary somatosensory area, mouth, layer 4 | Subparafascicular nucleus, magnocellular part |
| Primary somatosensory area, mouth, layer 5 | Subparafascicular nucleus, parvicellular part |
| Primary somatosensory area, mouth, layer 6a | Subparaventricular zone |
| Primary somatosensory area, mouth, layer 6b | Substantia innominata |
| Primary somatosensory area, nose, layer 1 | Substantia nigra, compact part |
| Primary somatosensory area, nose, layer 2/3 | Substantia nigra, reticular part |
| Primary somatosensory area, nose, layer 4 | Subthalamic nucleus |
| Primary somatosensory area, nose, layer 5 | Superior colliculus, motor related, deep gray layer |
| Primary somatosensory area, nose, layer 6a | Superior colliculus, motor related, deep white layer |
| Primary somatosensory area, nose, layer 6b | Superior colliculus, motor related, intermediate gray layer |
| Primary somatosensory area, trunk, layer 1 | Superior colliculus, motor related, intermediate gray layer, sublayer a |
| Primary somatosensory area, trunk, layer 2/3 | Superior colliculus, motor related, intermediate gray layer, sublayer b |
| Primary somatosensory area, trunk, layer 4 | Superior colliculus, motor related, intermediate gray layer, sublayer c |
| Primary somatosensory area, trunk, layer 5 | Superior colliculus, motor related, intermediate white layer |
| Primary somatosensory area, trunk, layer 6a | Superior colliculus, optic layer |
| Primary somatosensory area, trunk, layer 6b | Superior colliculus, superficial gray layer |
| Primary somatosensory area, unassigned, layer 2/3 | Supplemental somatosensory area, layer 1 |
| Primary somatosensory area, unassigned, layer 4/5/6a/6b | Supplemental somatosensory area, layer 2/3 |
| Primary somatosensory area, upper limb, layer 1 | Supplemental somatosensory area, layer 4 |
| Primary somatosensory area, upper limb, layer 2/3 | Supplemental somatosensory area, layer 5 |
| Primary somatosensory area, upper limb, layer 4 | Supplemental somatosensory area, layer 6a |
| Primary somatosensory area, upper limb, layer 5 | Supplemental somatosensory area, layer 6b |
| Primary somatosensory area, upper limb, layer 6a | Suprachiasmatic nucleus |
| Primary somatosensory area, upper limb, layer 6b | Suprageniculate nucleus |
| Primary visual area, layer 1 | Supramammillary nucleus |
| Primary visual area, layer 2/3 | Supraoptic nucleus |
| Primary visual area, layer 4 | Taenia tecta |
| Primary visual area, layer 5 | Temporal association areas, layer 1 |
| Primary visual area, layer 6a | Temporal association areas, layer 2/3 |
| Primary visual area, layer 6b | Temporal association areas, layer 4 |
| Pyramus (VIII) | Temporal association areas, layer 5 |
| Red nucleus | Temporal association areas, layer 6a |
| Reticular nucleus of the thalamus | Temporal association areas, layer 6b |
| Retrochiasmatic area | Thalamus |
| Retrosplenial area, dorsal part, layer 1 | Triangular nucleus of septum |
| Retrosplenial area, dorsal part, layer 2/3 | Trochlear nucleus |
| Retrosplenial area, dorsal part, layer 4 | Tuberal nucleus |
| Retrosplenial area, dorsal part, layer 5 | Tuberomammillary nucleus, dorsal part |
| Retrosplenial area, dorsal part, layer 6a | Tuberomammillary nucleus, ventral part |
| Retrosplenial area, dorsal part, layer 6b | Uvula (IX) |
| Retrosplenial area, lateral agranular part | Vascular organ of the lamina terminalis |
| Retrosplenial area, lateral agranular part, layer 1 | Ventral anterior-lateral complex of the thalamus |
| Retrosplenial area, lateral agranular part, layer 2/3 | Ventral auditory area, layer 1 |
| Retrosplenial area, lateral agranular part, layer 5 | Ventral auditory area, layer 2/3 |
| Retrosplenial area, lateral agranular part, layer 6a | Ventral auditory area, layer 4 |
| Retrosplenial area, ventral part, layer 1 | Ventral auditory area, layer 5 |
| Retrosplenial area, ventral part, layer 2/3 | Ventral auditory area, layer 6a |
| Retrosplenial area, ventral part, layer 5 | Ventral auditory area, layer 6b |
| Retrosplenial area, ventral part, layer 6a | Ventral medial nucleus of the thalamus |
| Retrosplenial area, ventral part, layer 6b | Ventral part of the lateral geniculate complex |
| Rhomboid nucleus | Ventral posterolateral nucleus of the thalamus |
| Rostral linear nucleus raphe | Ventral posterolateral nucleus of the thalamus, parvicellular part |
| Rostrolateral visual area | Ventral posteromedial nucleus of the thalamus |
| Secondary motor area, layer 1 | Ventral posteromedial nucleus of the thalamus, parvicellular part |
| Secondary motor area, layer 2/3 | Ventral premammillary nucleus |
| Secondary motor area, layer 5 | Ventral tegmental area |
| Secondary motor area, layer 6a | Ventral tegmental nucleus |
| Secondary motor area, layer 6b | Ventrolateral preoptic nucleus |
| Septofimbrial nucleus | Ventromedial hypothalamic nucleus |
| Septohippocampal nucleus | Visceral area, layer 1 |
| Simple lobule | Visceral area, layer 2/3 |
| Striatum | Visceral area, layer 4 |
| Striatum-like amygdalar nuclei | Visceral area, layer 5 |
| Subfornical organ | Visceral area, layer 6a |
| Subgeniculate nucleus | Visceral area, layer 6b |
| Subiculum | Zona incerta |

### Table S2. List of 150 brain regions and their individual layers with corresponding engram indices, Related to Figure 2

| Brain regions | Engram index |
|---|---|
| Anterodorsal nucleus | 2.923 |
| Pallidum | 1.889 |
| Dentate gyrus, polymorph layer | 1.715 |
| Dentate gyrus, molecular layer | 1.359 |
| Superior colliculus, motor related, deep white layer | 1.266 |
| Periaqueductal gray | 1.096 |
| Flocculus | 1.026 |
| Fastigial nucleus | 1.010 |
| Superior colliculus, motor related, intermediate white layer | 1.002 |
| Ventral posteromedial nucleus of the thalamus | 0.822 |
| Posterolateral visual area, layer 4 | 0.765 |
| Ventral auditory area, layer 4 | 0.749 |
| Superior colliculus, optic layer | 0.743 |
| Ventral medial nucleus of the thalamus | 0.734 |
| Caudoputamen | 0.682 |
| Infralimbic area, layer 1 | 0.667 |
| Paraventricular hypothalamic nucleus | 0.645 |
| Thalamus | 0.625 |
| Dorsal auditory area, layer 1 | 0.595 |
| Dorsal part of the lateral geniculate complex | 0.586 |
| Ventral auditory area, layer 1 | 0.564 |
| Red nucleus | 0.554 |
| Gustatory areas, layer 2/3 | 0.549 |
| Agranular insular area, posterior part, layer 2/3 | 0.524 |
| Temporal association areas, layer 4 | 0.519 |
| Postpiriform transition area | 0.514 |
| Field CA2 | 0.488 |
| Paraflocculus | 0.481 |
| Agranular insular area, dorsal part, layer 1 | 0.473 |
| Basolateral amygdalar nucleus, ventral part | 0.448 |
| Substantia innominata | 0.441 |
| Superior colliculus, superficial gray layer | 0.439 |
| Midbrain trigeminal nucleus | 0.434 |
| Secondary motor area, layer 1 | 0.430 |
| Cortical amygdalar area, anterior part | 0.424 |
| Nucleus of the lateral olfactory tract, layer 3 | 0.419 |
| Primary auditory area, layer 2/3 | 0.407 |
| Posterior hypothalamic nucleus | 0.401 |
| Striatum | 0.396 |
| Inferior colliculus | 0.395 |
| Posterior complex of the thalamus | 0.393 |
| Substantia nigra, reticular part | 0.391 |
| Central lateral nucleus of the thalamus | 0.390 |
| Reticular nucleus of the thalamus | 0.385 |
| Field CA3 | 0.383 |
| Agranular insular area, dorsal part, layer 2/3 | 0.382 |
| Posterolateral visual area, layer 5 | 0.370 |
| Dentate gyrus, granule cell layer | 0.369 |
| Gustatory areas, layer 5 | 0.365 |
| Lateral dorsal nucleus of thalamus | 0.364 |
| Dentate nucleus | 0.344 |
| Central amygdalar nucleus | 0.327 |
| Midbrain reticular nucleus | 0.319 |
| Parafascicular nucleus | 0.307 |
| Ventral auditory area, layer 2/3 | 0.303 |
| Nucleus of the posterior commissure | 0.298 |
| Anteromedial nucleus | 0.286 |
| Nucleus accumbens | 0.285 |
| Anterior pretectal nucleus | 0.285 |
| Basomedial amygdalar nucleus | 0.281 |
| Presubiculum | 0.265 |
| Temporal association areas, layer 2/3 | 0.264 |
| Endopiriform nucleus, ventral part | 0.261 |
| Field CA1 | 0.257 |
| Retrosplenial area, dorsal part, layer 4 | 0.251 |
| Ectorhinal area/Layer 1 | 0.248 |
| Primary motor area, Layer 2/3 | 0.247 |
| Primary auditory area, layer 6b | 0.237 |
| Midbrain | 0.235 |
| Frontal pole, layer 2/3 | 0.234 |
| Suprageniculate nucleus | 0.227 |
| Ventral auditory area, layer 5 | 0.212 |
| Visceral area, layer 5 | 0.209 |
| Temporal association areas, layer 1 | 0.201 |
| Orbital area, lateral part, layer 1 | 0.200 |

| Brain regions (continued) | Engram index |
|---|---|
| Taenia tecta | 0.191 |
| Dorsal peduncular area | 0.190 |
| Basolateral amygdalar nucleus, posterior part | 0.189 |
| Frontal pole, layer 1 | 0.186 |
| Piriform-amygdalar area | 0.184 |
| Olfactory tubercle | 0.182 |
| Agranular insular area, posterior part, layer 5 | 0.180 |
| Temporal association areas, layer 5 | 0.177 |
| Primary auditory area, layer 5 | 0.177 |
| Ectorhinal area/Layer 2/3 | 0.173 |
| Posterior auditory area, layer 2/3 | 0.171 |
| Intercalated amygdalar nucleus | 0.165 |
| Medial amygdalar nucleus | 0.164 |
| Lateral amygdalar nucleus | 0.157 |
| Primary somatosensory area, upper limb, layer 2/3 | 0.157 |
| Agranular insular area, ventral part, layer 2/3 | 0.153 |
| Cortical amygdalar area, posterior part | 0.150 |
| Primary auditory area, layer 6a | 0.144 |
| Medial geniculate complex | 0.142 |
| Primary auditory area, layer 4 | 0.142 |
| Interposed nucleus | 0.139 |
| Claustrum | 0.134 |
| Fields of Forel | 0.133 |
| Retrosplenial area, lateral agranular part, layer 1 | 0.128 |
| Medial pretectal area | 0.127 |
| Cuneiform nucleus | 0.125 |
| Lateral visual area, layer 6a | 0.118 |
| Mediodorsal nucleus of thalamus | 0.115 |
| Septofimbrial nucleus | 0.114 |
| Zona incerta | 0.107 |
| Agranular insular area, posterior part, layer 6a | 0.104 |
| Perirhinal area, layer 5 | 0.103 |
| Basolateral amygdalar nucleus, anterior part | 0.103 |
| Magnocellular nucleus | 0.101 |
| Ectorhinal area/Layer 5 | 0.098 |
| Lateral visual area, layer 2/3 | 0.098 |
| Olfactory areas | 0.097 |
| Postsubiculum | 0.096 |
| Ventral auditory area, layer 6a | 0.093 |
| Substantia nigra, compact part | 0.088 |
| Entorhinal area, lateral part | 0.086 |
| Lateral habenula | 0.083 |
| Nucleus of reunions | 0.080 |
| Cortical subplate | 0.075 |
| Agranular insular area, ventral part, layer 5 | 0.073 |
| Gustatory areas, layer 6a | 0.068 |
| Posterior limiting nucleus of the thalamus | 0.055 |
| Subiculum | 0.052 |
| Entorhinal area, medial part, dorsal zone | 0.049 |
| Striatum-like amygdalar nuclei | 0.036 |
| Primary somatosensory area, unassigned, layer 2/3 | 0.032 |
| Posterior amygdalar nucleus | 0.030 |
| Dorsal auditory area, layer 2/3 | 0.029 |
| Primary somatosensory area, upper limb, layer 4 | 0.015 |
| Temporal association areas, layer 6a | 0.009 |
| Primary motor area, Layer 5 | 0.004 |
| Lateral visual area, layer 4 | -0.013 |
| Secondary motor area, layer 2/3 | -0.029 |
| Laterointermediate area | -0.039 |
| Ectorhinal area/Layer 6a | -0.050 |
| Dorsal auditory area, layer 4 | -0.050 |
| Endopiriform nucleus, dorsal part | -0.051 |
| Primary somatosensory area, upper limb, layer 5 | -0.057 |
| Lateral septal nucleus, rostral (rostroventral) part | -0.057 |
| Primary somatosensory area, lower limb, layer 5 | -0.077 |
| Lateral visual area, layer 5 | -0.091 |
| Primary visual area, layer 2/3 | -0.095 |
| Primary visual area, layer 4 | -0.103 |
| Primary somatosensory area, lower limb, layer 2/3 | -0.103 |
| Dorsal auditory area, layer 6a | -0.105 |
| Primary visual area, layer 6a | -0.111 |
| Anterolateral visual area, layer 6a | -0.131 |
| Dorsal auditory area, layer 5 | -0.144 |
| Primary somatosensory area, lower limb, layer 4 | -0.163 |
| Hippocampal formation | -0.176 |

#### CONTACT FOR REAGENT AND RESOURCE SHARING

#### EXPERIMENTAL MODEL AND SUBJECT DETAILS

##### Animals

The C57BL/6J wild type male mice were obtained from Jackson Laboratory. For brain-wide neural activity labeling based on the *c-fos* promoter, we used the previously described c-fos-Cre$^{ERT2}$ mouse line (Guenthner et al., 2013). These mice are also known as Fos$^{CreER}$ or FosTRAP mice in which cFos-positive neurons can be labeled by the intraperitoneal injection of 4-hydroxytamoxifen (4-OHT) within a user-defined time-window. For our brain-wide labeling experiments, FosTRAP mice were crossed with the Cre-dependent tdTomato reporter mouse line Ai14, which were obtained from Jackson Laboratory (Stock No. 007908). All mouse lines were maintained as hemizygotes. Mice had access to food and water *ad libitum* and were socially housed in numbers of two to five littermates until surgery. Following surgery, mice were singly housed. For behavioral experiments, all mice were male and 3-5 months old. For virus-mediated activity-dependent labeling experiments (Roy et al., 2016), wild type male mice had been raised on food containing 40 mg kg$^{-1}$ doxycycline (DOX) for at least one week before surgery and remained on DOX for the remainder of the experiments except for 24 hours preceding the target-labeling day. For brain-wide activity dependent labeling followed by SHIELD tissue clearing of different behavioral epochs, male mice were 3-6 months old at the time of 4-OHT labeling. All experiments were conducted in accordance with U.S. National Institutes of Health (NIH) guidelines and the Massachusetts Institute of Technology Department of Comparative Medicine and Committee of Animal Care.

#### METHOD DETAILS

##### Brain-wide activity-dependent labeling

For brain-wide labeling experiments, FosTRAP mice crossed to Ai14 reporter mice were employed. 4-OHT (Sigma-Aldrich) was dissolved in 100% ethanol solution by shaking at 37°C for 20-30 min. One-part castor oil to four parts sunflower oil was combined to prepare the oil mixture that would eventually be injected intraperitoneally (IP) into the mouse. Dissolved 4-OHT was combined with the oil mixture, followed by ethanol evaporation using a centrifuge. The final concentration of 4-OHT dissolved in the oil mixture was 10 mg ml$^{-1}$. For each male mouse, optimal activity-dependent labeling was achieved using a target concentration of 30-40 mg kg$^{-1}$. One hour prior to the behavioral epoch of interest, mice were injected with 4-OHT. Following behavior experiments, mice were returned to their home cages and remained undisturbed for at least 72 hours.

##### SHIELD processing and clearing

SHIELD tissue processing was performed as previously described (Park et al., 2018). Briefly, mice were perfused first with ice-cold 1x PBS solution, followed by ice-cold SHIELD perfusion solution (4% w/v paraformaldehyde with the supernatant of 10% w/v polyglycerol 3-polyglycidyl ether resin prepared in 1x PBS). Polyglycerol 3-polyglycidyl ether (P3PE) resin was provided by

EPM-CVC Thermoset Specialties. After 2 days of incubation in the perfusion solution at 4°C, each brain sample was split into two hemispheres and further incubated in the supernatant of 10% w/v P3PE resin prepared in 1x PBS at 4°C for 24 hours. After 24 hours of subsequent incubation in 0.1 M carbonate buffer solution (pH 10.0) at 37°C, brain hemispheres were transferred to 1x PBS solution containing 0.02% sodium azide and were stored until the delipidation process. For delipidation, brain hemispheres were incubated in a solution containing 10 mM sodium borate, 100 mM sodium sulfite, and 300 mM sodium dodecyl sulfate (pH 9.0 using sodium hydroxide) at 37°C for 1 day, followed by 8~10 days of incubation at 45°C with shaking. Once the brain hemispheres were rendered evenly translucent based on visual inspection, they were incubated in iohexol-based PROTOS solution (125 g iohexol, 3 g diatrizoic acid, and 5 g N-methyl-D-glucamine in ~110 ml distilled water with a final refractive index set to 1.458) for over 24 hours at room temperature for optical clearing.

**Light-sheet microscope imaging**

Transparent brain hemispheres were secured onto a sample holder using 1.5% agarose prepared in the PROTOS solution. After the tissue-agarose mold was fully equilibrated in the PROTOS solution (no visible haze at the tissue-agarose interface and the agarose-solution interface), individual brain hemispheres were imaged using a custom-built light-sheet microscope. Brain-wide tdTomato signal detection (excited by a 561 nm laser) and autofluorescence signal detection (excited by a 635 nm laser) was performed using a 10x/0.6NA objective lens. After image acquisition, datasets were 4x down-sampled in the xy plane to obtain 2.34 x 2.34 x 5 µm (x, y, z) voxel size. Illumination correction was performed with a custom-generated MATLAB script, and images were stitched using TeraStitcher.

**Quantitative activity mapping**

Brain hemisphere autofluorescence images were down-sampled to 25 x 25 x 25 µm voxel size and automatically aligned to the annotated autofluorescence atlas from the Allen Brain Institute (version 3) (Oh et al., 2014) by linear- and non-linear image transformation processes. After manually validating alignment accuracy for individual samples, tdTomato images were projected in order to perform spot detection by local maxima and watershed transformation. Refinement of detected spots was done through a neural net model built from a manual classification of ~10,000 cells. Even though the trained neural net showed 91% accuracy (95.7% true-negative and 85.8% true-positive rates), detection accuracy of individual hemisphere datasets was confirmed by manual inspection. Detected spot information combined with atlas alignment data enabled the quantification of brain region-specific tdTomato$^{+}$ cell counts. Activated neuron counts were obtained from 409 individual brain regions from each hemisphere except the medulla, because the discrepancy between autofluorescence from this structure in cleared brain hemispheres vs. version 3 of the Allen Brain reference atlas (generated from PFA-fixed brain sections) interfered with automatic region segmentation of this structure. We also excluded fiber tracts since the number of activated neurons in these structures was negligible.

Cell counts from cohorts of behavioral groups, specifically context (Ctx), CFC, recall (Re), were normalized to average counts from the home cage (HC) group. The engram index was calculated for each brain region using the following equation:

$$Engram\ index = \log_{10}\left(\frac{\overline{\mu}_{CFC} - \overline{\mu}_{Ctx}}{\overline{\mu}_{CFC} - \overline{\mu}_{Re}}\right)$$

where $\mu$ is the mean number of activated neurons for individual behavioral groups. This equation includes a common logarithm (base 10) for plotting purposes only. The engram index was only calculated for brain regions in which CFC-Recall group counts were significantly higher than that of Home Cage-Context groups (two-tailed *t* test, $P < 0.05$). Imaris software was used for 3D rendering of tdTomato hemisphere images. Data plots and heat map generation were performed using custom MATLAB scripts.

Figure 1B shows a representative heat map for the brain-wide activity mapping experiments. All brain regions, including their sub-layers, that were quantified in these activity mapping experiments are listed in Table S1. Figures 2B and 2C shows a rank-ordered list of brain regions with a heat map of engram index values from high to low (i.e., engram index values greater than 0). These regions are the ones in which CFC-Recall group counts were significantly higher than that of Home Cage-Context groups. For the regions in Figures 2B and 2C that have sub-layers, an expanded list of regions and their individual layers along with engram index values are provided in Table S2.

**Viral constructs**

To label memory engram cells in wild type mice maintained on DOX food, we used a double-virus system that combined the c-Fos-tTA virus with a TRE-dependent virus. The pAAV-c-Fos-tTA plasmid was previously described (Roy et al., 2016). Similarly, the following TRE-dependent constructs were also previously described (Liu et al., 2012; Ryan et al., 2015): pAAV-TRE-ChR2-eYFP, pAAV-TRE-eYFP, pAAV-TRE-mCherry. The pAAV-TRE-eArchT-mCherry and pAAV-TRE-hM3Dq-mCherry plasmids were constructed by introducing the eArchT and hM3Dq fragments respectively, into the pAAV-TRE-mCherry plasmid backbone. AAV vectors were serotyped with $AAV_9$ coat proteins and packaged at the University of Massachusetts Medical School Gene Therapy Center and Vector Core, or Vigene Biosciences. Viral titers were $1.5 \times 10^{13}$ genome copy (GC) $ml^{-1}$ for $AAV_9$-c-Fos-tTA, $AAV_9$-TRE-ChR2-eYFP and $AAV_9$-TRE-eYFP, $2 \times 10^{13}$ GC $ml^{-1}$ for $AAV_9$-TRE-eArchT-mCherry, $3 \times 10^{13}$ GC $ml^{-1}$ for $AAV_9$-TRE-mCherry, and $1.3 \times 10^{13}$ GC $ml^{-1}$ for $AAV_9$-TRE-hM3Dq-mCherry.

**Surgery and optic fiber implants**

Mice were anesthetized with isoflurane or 500 mg $kg^{-1}$ avertin for stereotaxic injections. Injections were targeted bilaterally to primary visual cortex (V1; -2.7 mm AP, +/-2.5 mm ML, -1.1 mm DV), primary somatosensory cortex (S1BF; -1.58 mm AP, +/-2.75 mm ML, -1.5 mm DV), basolateral amygdala (BLA; -1.46 mm AP, +/-3.3 mm ML, -4.68 mm DV), anteromedial thalamic nucleus (AM; -0.7 mm AP, +/-0.63 mm ML, -3.7 mm DV), dorsal CA1 (-2.1 mm AP, +/-1.5 mm ML, -1.4 mm DV), and thalamic nucleus reuniens (RE; -0.58 mm AP, +/-0.25 mm ML, -4.15 mm DV). Injection volumes were 300 nl for V1 and S1BF, 200 nl for BLA, 150 nl for AM and RE, and 375 nl for CA1. Viruses were injected at 70 nl $min^{-1}$ using a glass micropipette attached to a 10 ml Hamilton microsyringe. The needle was lowered to the target site and remained for 5 min before beginning the injection. After the injection, the needle stayed for 10 min before it was withdrawn. Custom implants containing two optic fibers (200 mm core diameter; Doric Lenses) was lowered above the injection site for CA1 (0.15 mm above the injection DV). Single optic fiber implants

(200 mm core diameter; Doric Lenses) were lowered above the V1, S1BF, BLA, AM, and RE injection sites (0.15 mm above the injection DV). The implant was secured to the skull with two jewelry screws, adhesive cement (C&B Metabond) and dental cement. An opaque cap derived from the top part of a black Eppendorf tube protected the implant. Mice were given 1.5 mg $kg^{-1}$ metacam as analgesic and allowed to recover for 2 weeks before behavioral experiments. All injection sites were verified histologically. As criteria, we only included mice with virus expression limited to the targeted regions.

**Immunohistochemistry**

Mice were dispatched using 750–1000 mg $kg^{-1}$ avertin and transcardially perfused with 1x PBS solution, followed by 4% paraformaldehyde (PFA). Brains were extracted and post fixed in 4% PFA at 4°C for 24 hours. Brains were transferred to 1x PBS and 50 μm coronal slices were prepared using a vibratome. For immunostaining, each slice was placed in PBS + 0.3% Triton X-100 (PBS-T), with 5% normal goat serum for 1 hour and then incubated with primary antibody at 4°C for 24 hours. Slices then underwent three wash steps for 10 min each in PBS, followed by a 2-hour incubation with secondary antibody at room temperature. After three more wash steps of 10 min each in PBS-T, slices were mounted using VECTASHIELD mounting medium on positively-charged glass slides. Antibodies used for staining were as follows: chicken anti-GFP (1:1000, Life Technologies) and anti-chicken Alexa-488, rabbit anti-RFP (1:1000, Rockland Inc.) and anti-rabbit Alexa-555, cFos was stained with rabbit anti-cFos (1:400, Santa Cruz) and anti-rabbit Alexa-555, and nuclei were stained with DAPI (1:3000, Sigma). For counter staining in Figure 5, brain sections were incubated with Neuro Trace Fluorescent Nissl Stain (1:100, Molecular Probes) in 1x PBS for 1 hour after secondary antibody staining.

**Behavior assays**

Experiments were conducted during the light cycle (7 am to 7 pm). Mice were randomly assigned to experimental groups for each experiment. Mice were habituated to investigator handling for 1-2 minutes on three consecutive days. Handling took place in the holding room where the mice were housed. Prior to each handling session, mice were transported by wheeled cart to and from the vicinity of behavior rooms to habituate them to the journey. For natural memory recall sessions, data were quantified using FreezeFrame software. Optogenetic manipulations interfered with motion detection, and therefore freezing behavior in these experiments were manually quantified. All behavior experiments were analyzed blind to experimental group. Unpaired student's *t* tests were used for independent group comparisons, with Welch's correction when group variances were significantly different. Following behavioral protocols, brain sections were prepared to confirm efficient viral labeling in target areas. Animals lacking adequate labeling were excluded prior to behavior quantification.

**Contextual fear conditioning**

Two distinct contexts were employed for the contextual fear-conditioning (CFC) paradigm. The conditioning context were 29 × 25 × 22 cm chambers with grid floors, dim white lighting, and scented with 1% acetic acid. The neutral context consisted of 30 × 25 × 33 cm chambers with white perspex floors, red lighting, and scented with 0.25% benzaldehyde. All mice were conditioned (180 sec exploration, one 0.75 mA shock of 2 sec duration at 180 sec, second 0.75 mA shock of 2 sec duration at 240 sec, 120 sec post-shock period), and natural memory recall tests (3 min) were performed one day later. Experiments showed no generalization in the neutral context. Floors of chambers were cleaned with quatricide before and between runs. Mice were transported to and from the experimental room in their home cages using a wheeled cart. The cart and cages remained in an anteroom to the experimental rooms during all behavioral experiments. For activity-dependent labeling, mice were either injected with 4-OHT one hour prior to behavior (for FosTRAP experiments) or kept on regular food without DOX for 24 hours prior to training (for TRE/tTA experiments using B6 mice). For B6 mice, when training or recall was complete, mice were switched back to food containing 40 mg $kg^{-1}$ DOX.

**Optogenetic manipulations**

For light-induced freezing behavior, a context distinct from the CFC training chamber (context A) was used. These were 30 × 25 × 33 cm chambers with perspex floors, square ceilings, white lighting, and scented with 0.25% benzaldehyde. Chamber ceilings were customized to hold a rotary joint (Doric Lenses) connected to two 0.32 m patch cords. All mice had patch cords fitted to the optic fiber implant prior to testing. Two mice were run simultaneously in two identical chambers. ChR2 was stimulated at 4 or 20 Hz (15 ms pulse width) using a 473 nm laser (10-15 mW), for the designated epochs. Testing sessions were 6 min in duration, consisting of two 3 min epochs, with the first as a light-off epoch and the second as a light-on epoch. At the end of 6 min, the mouse was detached and returned to its home cage. Floors of chambers were cleaned with quatricide before and between runs. For green light inhibition experiments, inhibition was performed during the entire memory recall duration (3 min) using a 561 nm laser (~12 mW, constant green light).

**Chemogenetic activation**

For chemogenetic activation experiments, we employed the excitatory DREADDs receptor hM3Dq. These receptors are activated by the ligand clozapine-N-oxide (CNO), which is injected intraperitoneally (IP) into the mouse. For behavior experiments following CFC training and engram cell labeling, 4 mg $kg^{-1}$ CNO was injected IP 50 min before placing the mice in a context distinct from the CFC training chamber. Freezing behavior was automatically quantified during a 3 min session.

#### QUANTIFICATION AND STATISTICAL ANALYSIS

**Brain-wide activity mapping data analysis**

Data were analyzed using Prism 6 software. For data plotted in Figure 3, statistical comparisons used a one-way ANOVA followed by Tukey multiple comparison post-hoc tests ($*P < 0.05$, $**P < 0.01$). Data are presented as mean values accompanied by SEM.

**Freezing behavioral analysis**

Data are presented as mean values accompanied by SEM. No statistical methods were used to predetermine sample sizes. Data analysis was performed blind to the conditions of the experiments. Data were analyzed using Microsoft Excel with the Statplus plug-in and Prism 6 software. Statistical comparisons used unpaired *t* tests ($P > 0.05$ NS, $*P < 0.05$, $**P < 0.01$, $***P < 0.001$). Statistical parameters including the exact value of n, precision measures (mean ± SEM), and statistical significance are reported in figure legends.

**Manual cFos cell counts following engram cell manipulations**

Brain slices containing target brain regions were selected based on Nissl staining compared with a standard brain atlas (Franklin and Paxinos, 3rd edition). These sections were used to count the number of $cFos^+$ cells within individual regions (Figure 5). The number of $cFos^+$ cells in a 1 $mm^2$ area were counted using ImageJ and MATLAB software. Data were analyzed using Microsoft Excel and Prism 6 software. Heat maps represent cFos activation levels in natural recall, CA1 or BLA engram cell activation, and CA1 or BLA engram cell inhibition, respectively. Specifically, context A-A (trained context-induced memory recall) vs. context A-B (neutral context) for natural recall ratios, ChR2-eYFP vs. eYFP for engram cell activation ratios, and eArchT-mCherry vs. mCherry for engram cell inhibition ratios. Red colored heat map regions indicate an increase in the number of $cFos^+$ neurons based on the *p*-value obtained by comparing individual control vs. natural recall or manipulation group data, whereas blue colored regions indicate a decrease in the number of $cFos^+$ neurons (Figures 5I and 5S). These statistical comparisons used unpaired *t* tests. For scatter plots (Figures 5J and 5T), the ratio of $cFos^+$ cell counts were obtained by normalization to individual control group data. Each dot on the plot represents the ratio of $cFos^+$ cell counts in individual brain regions, and r represents the Pearson's correlation coefficient.
